# SCRAPPY - a single cell rapid assay of proteome perturbation in yeast uncovers a joint role of aromatic amino acids and oxidative stress in the toxicity of lipophilic nucleoside analogs

**DOI:** 10.1101/2024.02.19.580949

**Authors:** Eslam Ghazy, Victoria A. Bidiuk, Fedor Ryabov, Olga V. Mitkevich, Olga B. Riabova, Yaroslav M. Stanishevskiy, Igor B. Levshin, Liudmila A. Alexandrova, Maxim V. Jasko, Dmitriy A. Makarov, Alexander A. Zhgun, Darya A. Avdanina, Anna A. Ermolyuk, Vitaly V. Kushnirov, Anna P. Egorova, Michael O. Agaphonov, Alexander I. Alexandrov

## Abstract

Assaying cellular responses to antimicrobial molecules is a path to understanding modes of action of potential drugs. This is often achieved via transcriptomics and proteomics, but simple and inexpensive methods for rapid characterization are lacking. To bridge this gap, we assayed changes in the abundance of a panel of 64 “sentinel” proteins fused to GFP in the yeast *Saccharomyces cerevisiae* using flow cytometry. This method produced expected patterns for classical antifungals and allowed inference of common mechanisms between known and novel compounds. Single-cell data also revealed diverging responses in mitochondrial protein abundance in response to thiazolidine antifungals, and perturbations of the cell cycle caused by various compounds. Finally, the method provided insight into the unknown mode of action of alkylated nucleosides, which can be used against fungi residing on works of art. These substances elevate levels of proteins involved in the biosynthesis of aromatic amino acids (AAA), as well as in oxidative stress. Furthermore, deficiencies of Trp and Tyr biosynthesis increased the efficacy of these compounds, while antioxidants reduced it. Most surprisingly, antioxidant effectiveness relied on AAA biosynthesis. Thus, our approach and its possible modifications for other microbes provides an easy and reliable platform for revealing modes of action of novel compounds.

## INTRODUCTION

Fungal infections kill in excess of 1.5 million people annually and emergence of drug-resistant fungi is a global health risk [1]. This is mainly due to the severely limited number of approved classes of antifungal drugs. The main drugs used against systemic mycoses are azoles, which target the lanosterol demethylase Erg11 and inhibit ergosterol biosynthesis; polyenes, which bind ergosterol directly in the membrane; echinocandins, which inhibit glucan-synthase and thus inhibit cell wall synthesis; and fluoro-cytosine, which inhibits both transcription and replication. Additionally, there is a wider number of antifungal compounds which are used topically or in industrial applications. An example of such compounds is the copper ionophore zinc pyrithione, which is used in anti-dandruff treatment and in non-medical applications. For this compound, the influx of copper (or zinc) seems to damage numerous Fe-S proteins [2, 3] Importantly, resistance to nearly all of the compounds used for treatment of systemic mycoses has been reported in all of the standard fungal pathogens and emerging pathogens such as *Candida glabrata* and *C. auris* show inherent resistance to these drugs [1]. There is therefore an urgent need to create novel antifungal compounds and decipher their mechanisms of action.

Novel chemicals that affect living organisms are under constant development, including potential pharmaceuticals and drug-like substances, as well as various inhibitors which can be used for scientific research and other purposes. Whether these chemicals were developed rationally, or found during non-targeted screens, their effective use and further development requires detailed understanding of the effects on living organisms and the mechanisms of how these effects are realized. Common approaches to obtain these data are the identification of mutants with resistance or sensitivity to a chemical (which is most suitable for toxic substances) [4] or analysis of the cellular response to chemicals, most often using transcriptomic and mass-spectrometry (MS) based proteomic data (which are applicable to both toxic and non-toxic substances).

Notably, transcriptomics and MS-based proteomics provide very rich data on the response of a cell to a chemical or other treatment involving the quantities of thousands of transcripts or proteins, as well as information on splicing or post-translational modifications. However, these data are often expensive to obtain and analyze, and thus they can be limited in the number of conditions that can be tested. These approaches also do not easily provide single cell data on the cellular response to various treatments, which limits the sensitivity, and also might prevent observation of important phenomena which occur only in a limited subgroup of cells in a sample population. For instance, a small subset of a population can be persistors [5], i.e. cells with increased tolerance to a drug, which might thus exhibit a distinct response. Also, fungi can form biofilms [6], in which cells are often present in different morphologies, which may also influence resistance to antifungal agents [7] and thus, possibly, response profiles as well. Lastly, observation of responses in single-cell mode can be especially important for compounds that can kill the cells being studied, such as antimicrobials. Non-secretory proteins can leak out of cells or be excreted during necrotic cell death [8] and thus measuring their intracellular level would not be informative.

Due to the long available and powerful tools of gene engineering, *S. cerevisiae* is the most thoroughly studied fungus, and is among of the most studied eukaryotic organisms. Thus, inferring putative mechanisms of action of various compounds, including antifungals, from mutant screening as well as transcriptomic and proteomic responses, as well as follow-up hypothesis testing, using available systematic mutant collections [9], is comparatively easy in this organism. Numerous examples of obtaining useful information on the mechanisms of action of various compounds with antifungal activity have been reported [10, 11].

Systematic collections of yeast expressing GFP-tagged proteins [12] are highly productive when used for flow cytometry and have been used for high-throughput flow cytometric studies of various phenomena, such as protein level noise [13], replicative aging [14], as well as more focused studies, such as that of the neighboring gene effect [15]. Thus, this tool is ideal for obtaining single-cell proteomic data. Notably, a recent study has reported the utility of using a limited set of “sentinel proteins” to study the cellular response to stress using targeted mass-spectrometry-based proteomics [16].

The approach we offer and test is based on using flow cytometry to assay changes in the level of a limited set of 64 proteins tagged with GFP, with one strain of yeast being used to assay changes in the level of each protein (Figure 1A). This array was used to assay the response to 12 compounds, including three compounds used in the treatment of systemic mycoses (fluconazole, voriconazole and 5-fluorocytosine), a copper ionophore related to zinc pyrithione, and tunicamycin, all of which have known mechanisms of action, as well as several compounds for which the modes of action are unknown. Our results show that the method, termed SCRAPPY (single cell rapid assay of proteome perturbation in yeast), can be used to rapidly and easily obtain a proteomic profile of the cellular response to a wide range of chemicals. The obtained data show both common responses to a wide range of compounds, as well as compound-specific changes in the proteome. Follow-up study of a novel profile for a group of compounds with an unknown mechanism of action, alkylated cytidines, demonstrated the involvement of aromatic amino acid biosynthesis and oxidative stress, which proved to be interconnected.

**Figure 1.**
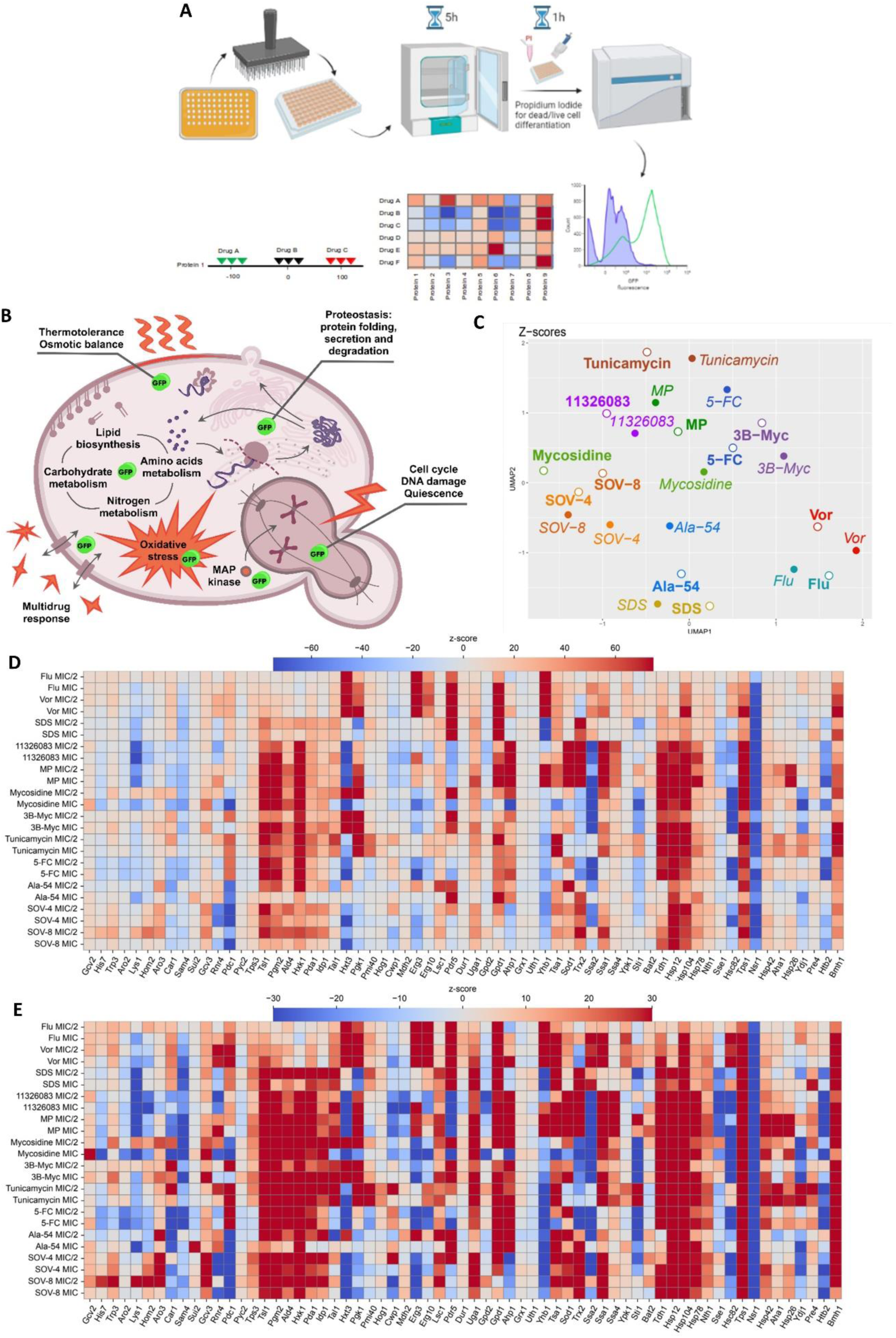
SCRAPPY provides clear profiles for various compounds, including classical antifungals, molecules with known targets and novel compounds. (A) Schematic of the SCRAPPY method and data outputs; (B) Schematic of the sentinel proteins included into the panel ; (C) UMAP representation of distances between averaged Z-score profiles for the tested compounds; (D) Changes in GFP-fusion protein abundance, using a Z-score metric with a −70/70 scale and (E) a −30/30 scale.

## MATERIAL AND METHODS

### Yeast strains and cell preparation for SCRAPPY

*Saccharomyces cerevisiae,* strain BY4741 (*MAT a his3Δ1 leu2Δ0 met15Δ0 ura3Δ0*) and its derivatives obtained in [16] were used in this work. For MIC determination, the wild type strain was inoculated into YPD medium (1% Yeast extract (w/v), 2% Peptone (w/v), 2% Glucose (w/v)) supplemented with 2-fold differing concentrations of tested compounds in 96 well plates (200 µL final volume), at an initial OD600 of 0.05. Cell growth was judged after 24 h of growth at 30^◦^C, with the MIC being the lowest concentration of drug where no growth was observed.

Derivatives of the BY4741 strain obtained in [12], harboring C-terminal GFP tags on proteins of interest, were used to detect changes of protein level in response to drug treatment. Genes encoding these proteins were modified in the genome at their 3’-terminus, which corresponds to the C-terminus of the protein, i.e., the regulatory regions of the genes were identical to those of the wild-type proteins. In general, this type of tagging rarely interferes with protein function or regulation[17]. A full table of the strains of *S. cerevisiae* used in this study is presented in Table 1. Cells bearing specific GFP fusions were pinned onto solid YPD medium (2% agar) (w/v), grown overnight to form small colonies, and then inoculated into YPD medium containing either no drug with appropriate DMSO concentration, or the tested drug at MIC or 0.5xMIC concentration, after which the cells were incubated for 6 h prior to flow cytometry. One hour prior to the end of incubation, propidium iodide was added to the incubating cells at a concentration of 1 µg/mL. After this, the control sample and drug-treated sample were analyzed on a Cytoflex S flow cytometer (Beckman Coulter) equipped with a 96-well sampler. GFP fluorescence was assayed using the 488 nm laser and 525/40 FITC filter, whereas propidium iodide staining was assayed with the 532 nm laser and 585/42 PE filter.

**Table 1.**
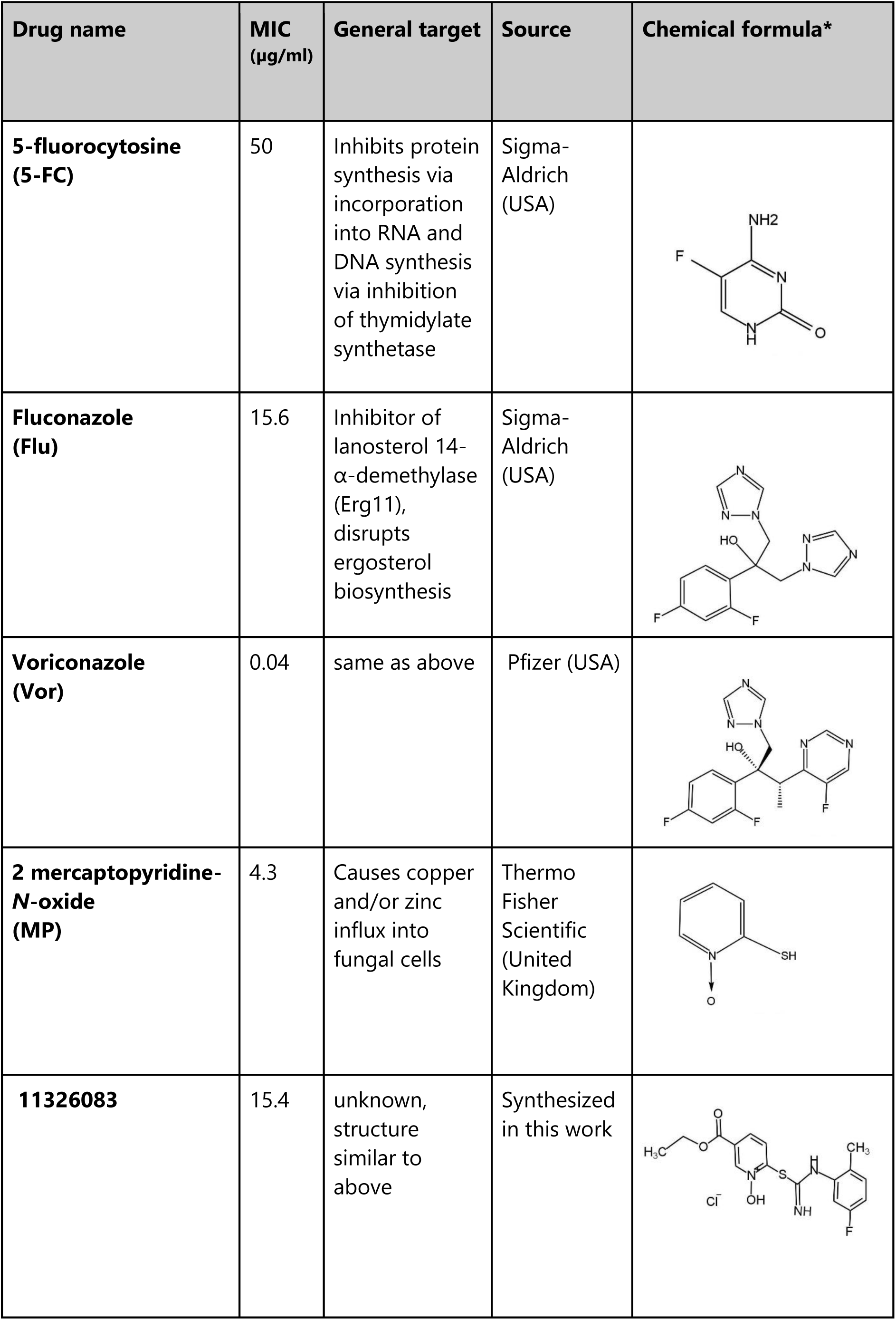
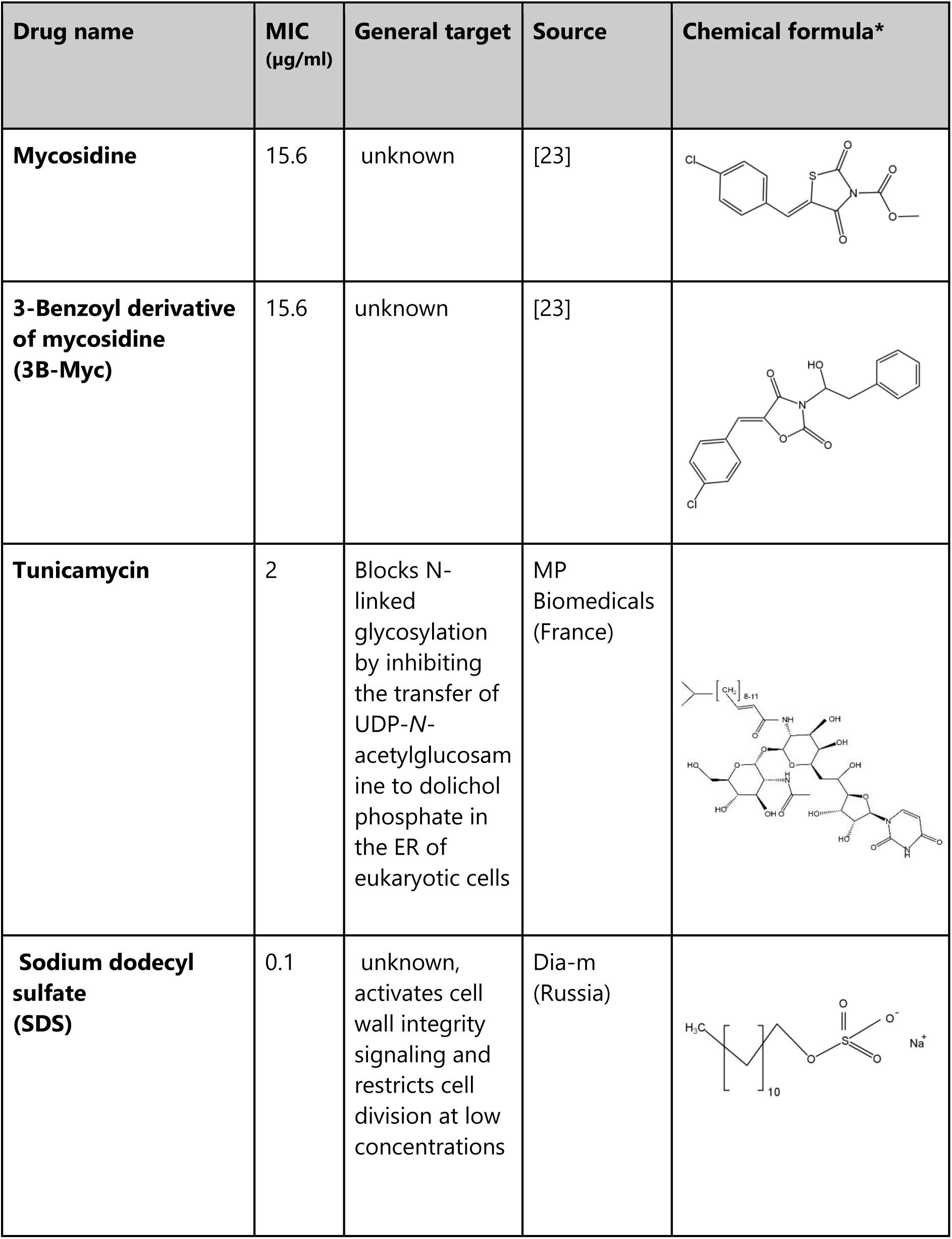
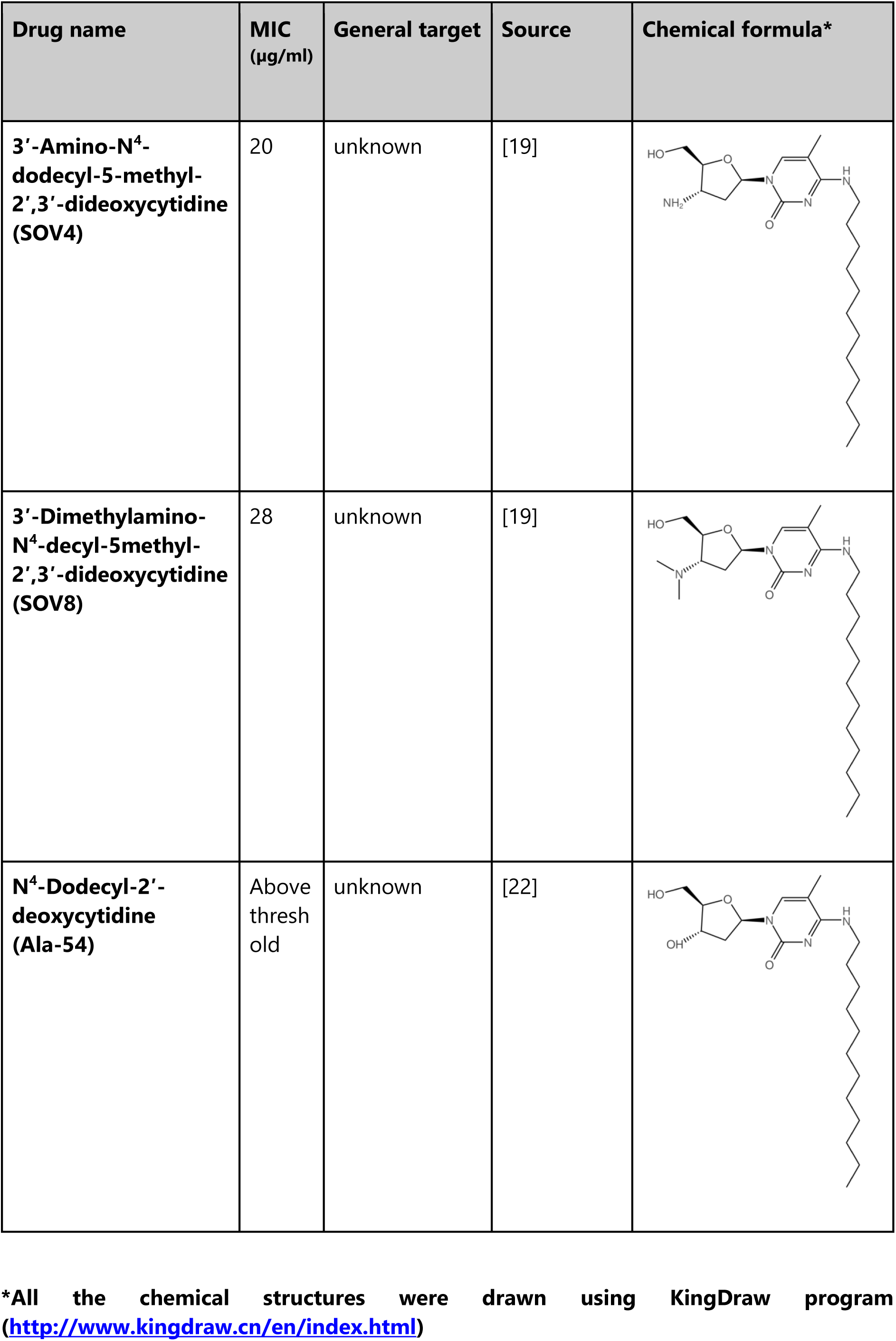
Chemical compounds used in this study.

| Drug name | MIC (µg/ml) | General target | Source | Chemical formula* |
| --- | --- | --- | --- | --- |
| **5-fluorocytosine (5-FC)** | 50 | Inhibits protein synthesis via incorporation into RNA and DNA synthesis via inhibition of thymidylate synthetase | Sigma-Aldrich (USA) |  |
| **Fluconazole (Flu)** | 15.6 | Inhibitor of lanosterol 14-α-demethylase (Erg11), disrupts ergosterol biosynthesis | Sigma-Aldrich (USA) |  |
| **Voriconazole (Vor)** | 0.04 | same as above | Pfizer (USA) |  |
| **2 mercaptopyridine-N-oxide (MP)** | 4.3 | Causes copper and/or zinc influx into fungal cells | Thermo Fisher Scientific (United Kingdom) |  |
| **11326083** | 15.4 | unknown, structure similar to above | Synthesized in this work |  |
| **Mycosidine** | 15.6 | unknown | [23] |  |
| **3-Benzoyl derivative of mycosidine (3B-Myc)** | 15.6 | unknown | [23] |  |
| **Tunicamycin** | 2 | Blocks N-linked glycosylation by inhibiting the transfer of UDP-N-acetylglucosamine to dolichol phosphate in the ER of eukaryotic cells | MP Biomedicals (France) |  |
| **Sodium dodecyl sulfate (SDS)** | 0.1 | unknown, activates cell wall integrity signaling and restricts cell division at low concentrations | Dia-m (Russia) |  |

| Drug name | MIC<br>(µg/ml) | General target | Source | Chemical formula* |
| --- | --- | --- | --- | --- |
| **3'-Amino-N<sup>4</sup>-dodecyl-5-methyl-2',3'-dideoxycytidine (SOV4)** | 20 | unknown | [19] |  |
| **3'-Dimethylamino-N<sup>4</sup>-decyl-5methyl-2',3'-dideoxycytidine (SOV8)** | 28 | unknown | [19] |  |
| **N<sup>4</sup>-Dodecyl-2'-deoxycytidine (Ala-54)** | Above thresh old | unknown | [22] |  |

\*All the chemical structures were drawn using KingDraw program (http://www.kingdraw.cn/en/index.html)

### Cultivation of Penicillium chrysogenum

*Penicillium chrysogenum* STG-117 (MW556011.1) was previously isolated from the surface of the icon ‘‘Prophet Solomon’’ (dated 1731) in the main historical building of the State Tretyakov Gallery (Moscow, Russia). Cultivation of this strain was carried out of the Czapek-Dox agar (CDA) medium under conditions as described previously [18, 19]. The toxic effect of N4-alkyl-cytidines (code names - SOVs and Ala-54) on the growth of fungal cells determined as previously described with some modifications [20] .. The inhibitory effect of the compounds was measured every 3 days for 48 days after inoculation from agar slants onto Petri dishes with CDA, supplemented with 0.2 mM SOV4, or SOV8, or Ala-54 or without additives (control). The percentage of fungal growth inhibition (FGI) was measured according to the formula: FGI % = [(Dc–Dt)/Dc] × 100, where Dc indicates the colony diameter in the control set and Dt indicates the colony diameter in the treatment set, as described in previous studies[19, 21]. The data recorded were measured in triplicate and repeated at least twice.

### Cytometry data analysis

Raw data was preliminarily analyzed using CytExpert software (Beckman Coulter). Then it was analyzed using custom python scripts (https://github.com/fedorrik/scrapper <u>and the</u> doi: 10.6084/m9.figshare.23736219). Comparison of protein levels between treated and control cells was performed using two metrics: fold change of fluorescence and z-score computation. For both options first the data obtained by dead cells was filtered (all data with PE-A > 5000 was removed). The protein level was calculated as the level of fluorescence divided by cell size (FITC-A / FSC-A) and the level of autofluorescence (FITC-A / FSC-A of a wild-type strain with no expressed GFP) subtracted. In case of fold change the median value of protein level of treated cells was divided by the median value of protein level of control cells.

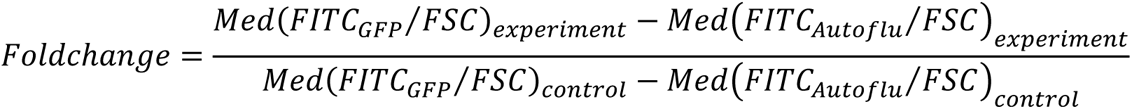

Z-scores were calculated as follows:

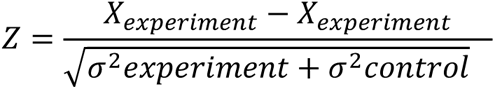

Where

- X _experiment_ = Med(FITC_GFP_ / FSC)_experiment_ - Med(FITC _Autoflu_ / FSC)_experiment_
- X _control_ = Med(FITC_GFP_ / FSC)_control_ - Med(FITC _Autoflu_ / FSC)_control_
- σ _experiment_ is the standard deviation of experiment sample divided by the square root of the number of data points
- σ _control_ is the standard deviation of control sample divided by the square root of the number of data points

The cytometry data has been deposited in http://flowrepository.org/,

### Synthesis of chemical substances

Table 2 presents the data on the compounds characterized in this study.

**Table 2.**
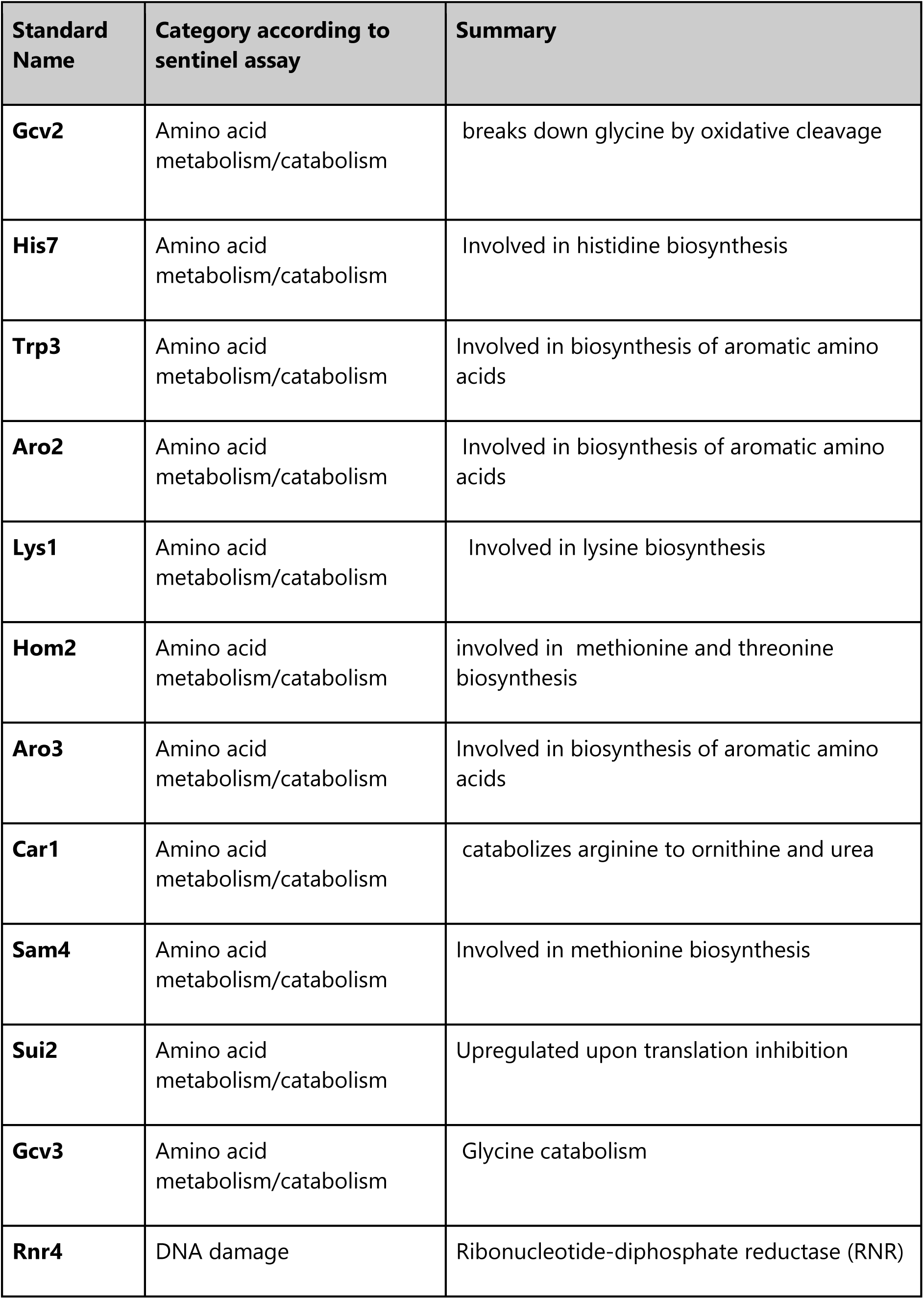
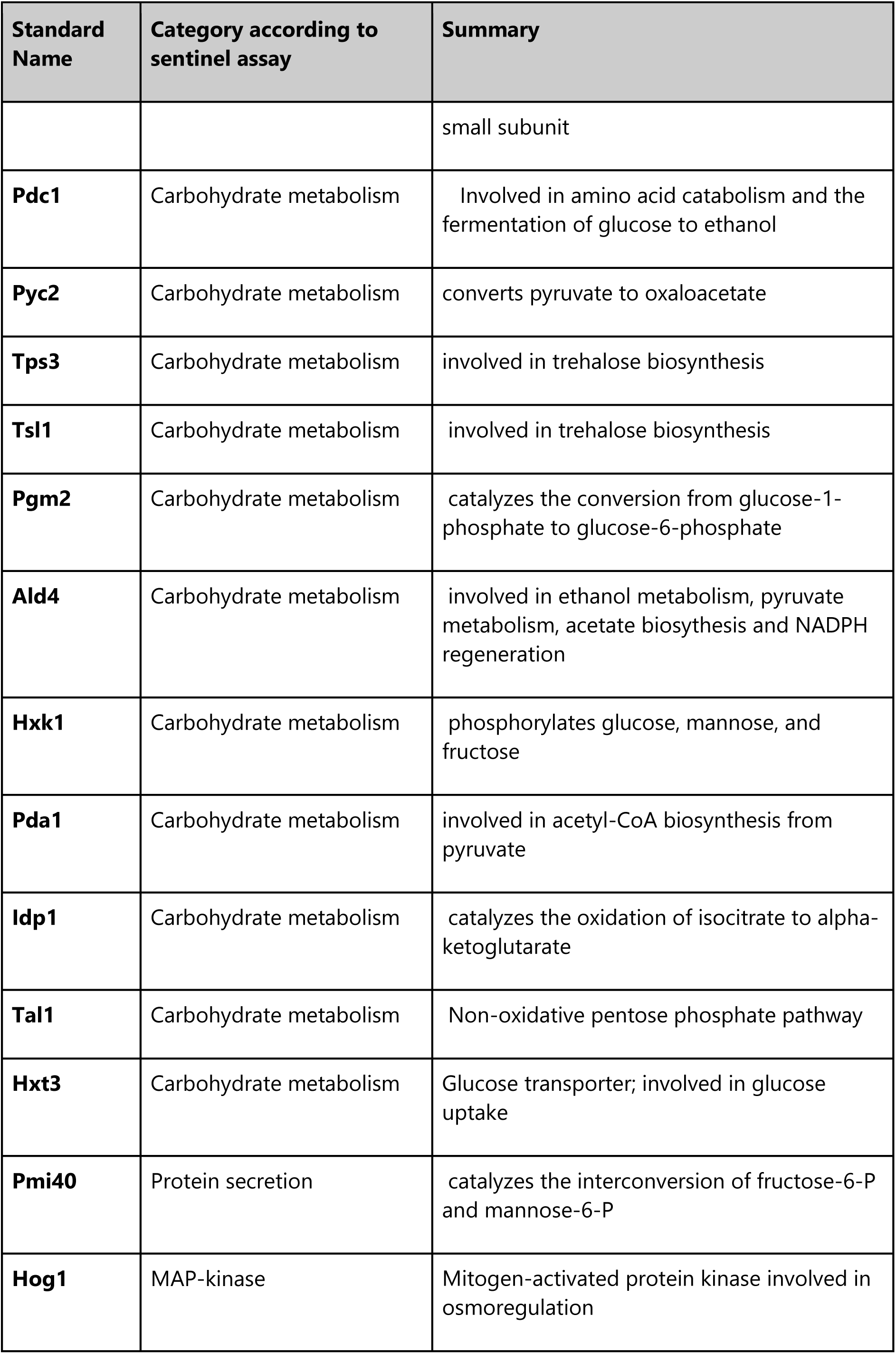
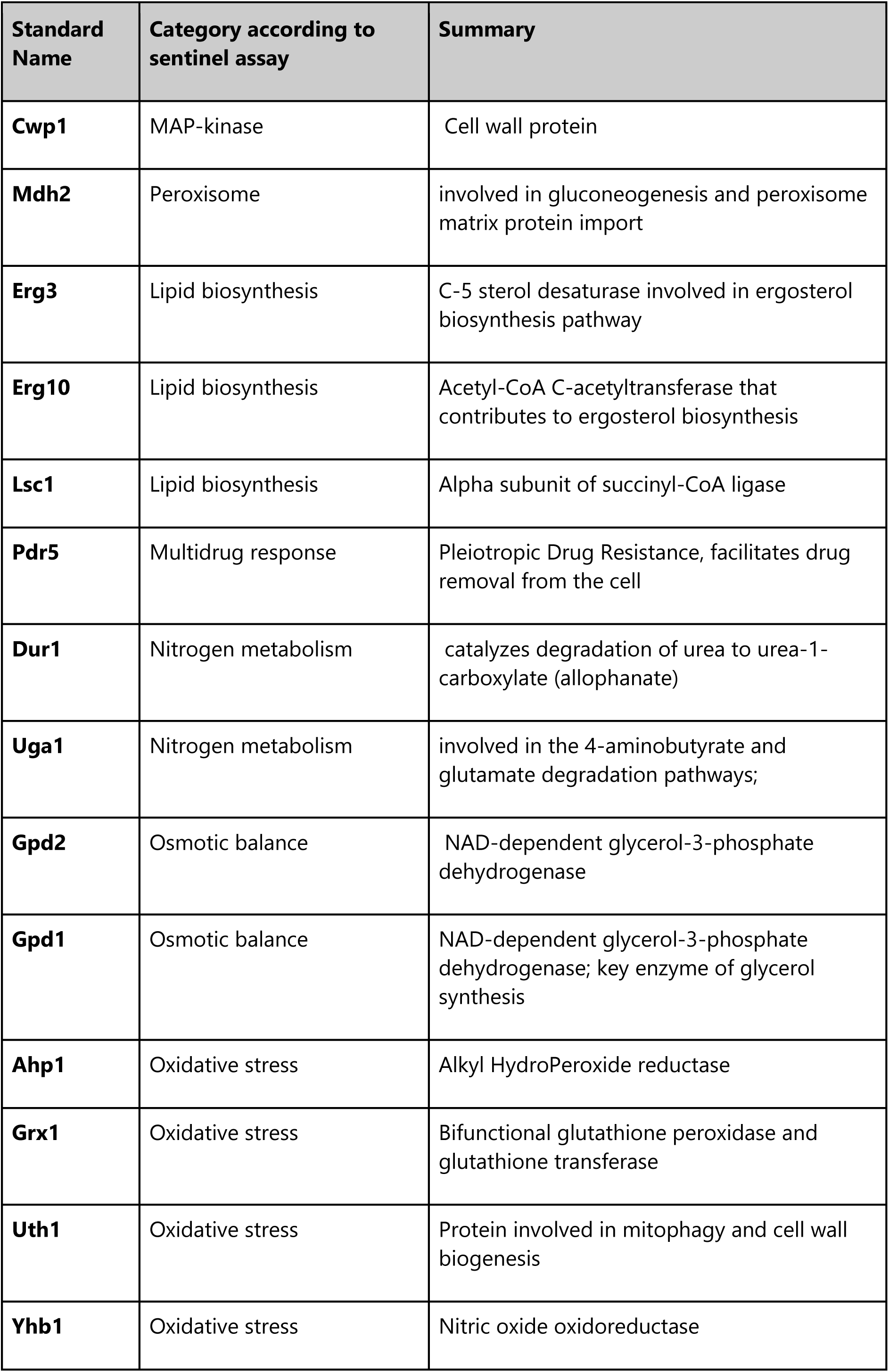
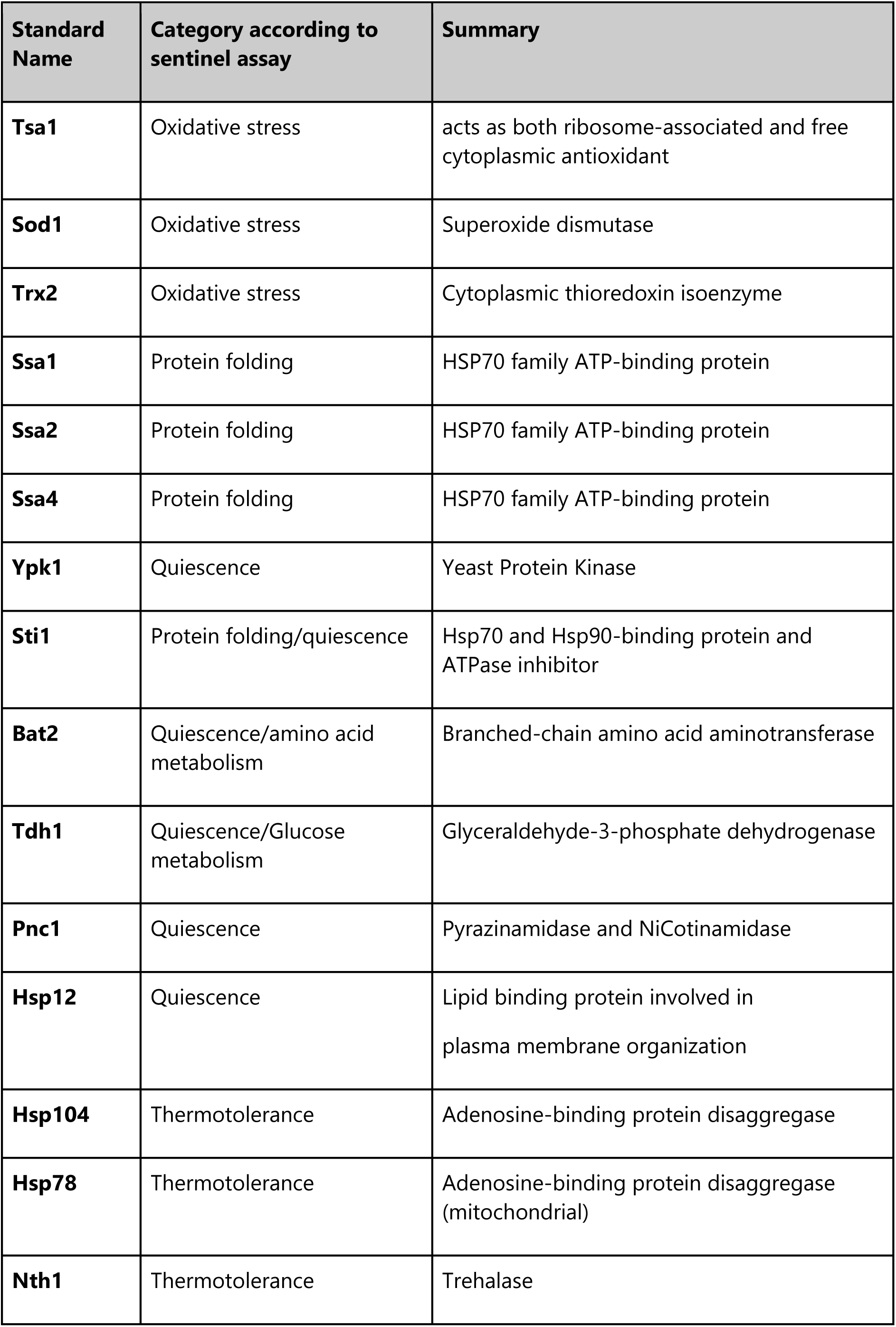
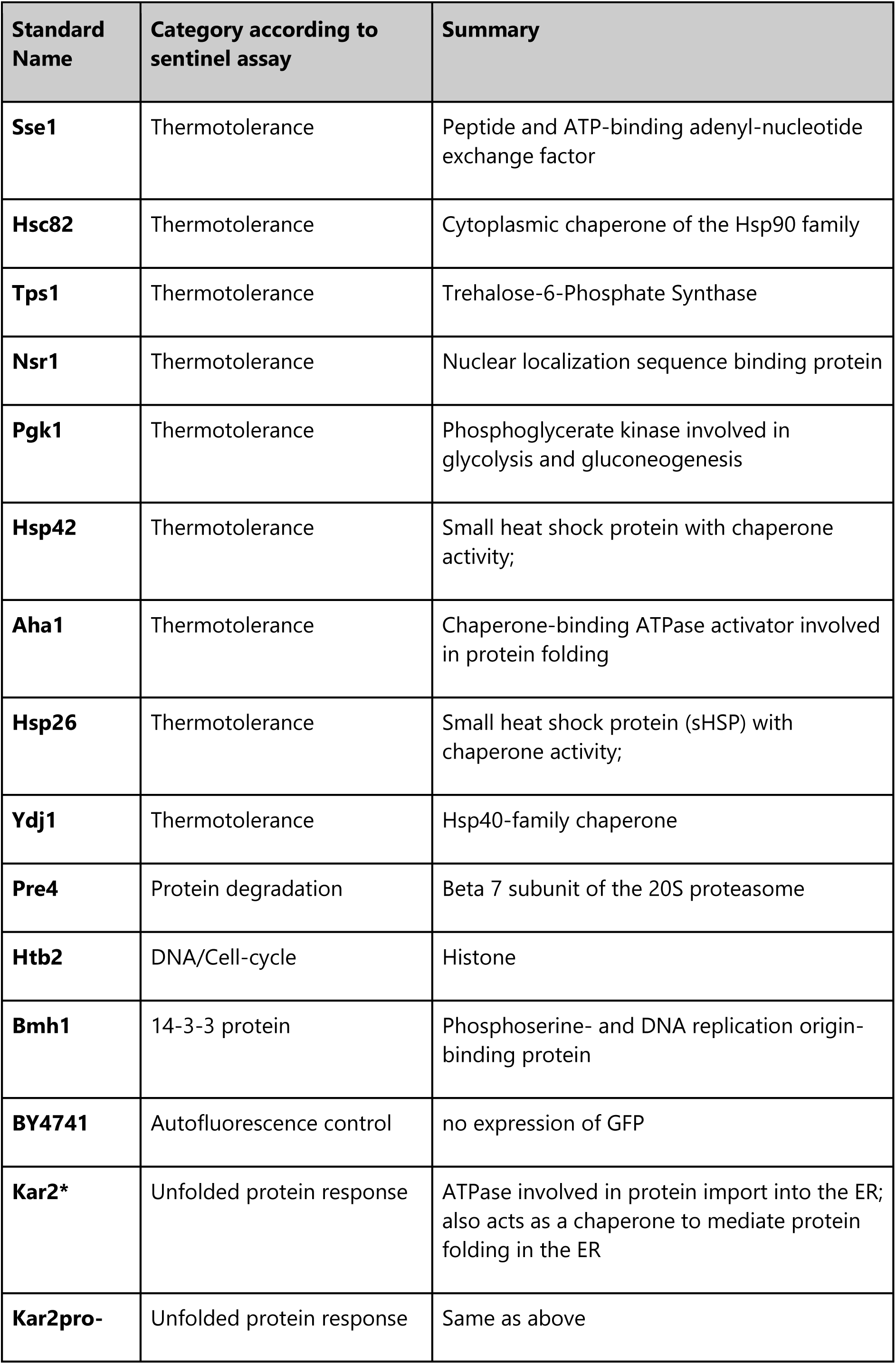
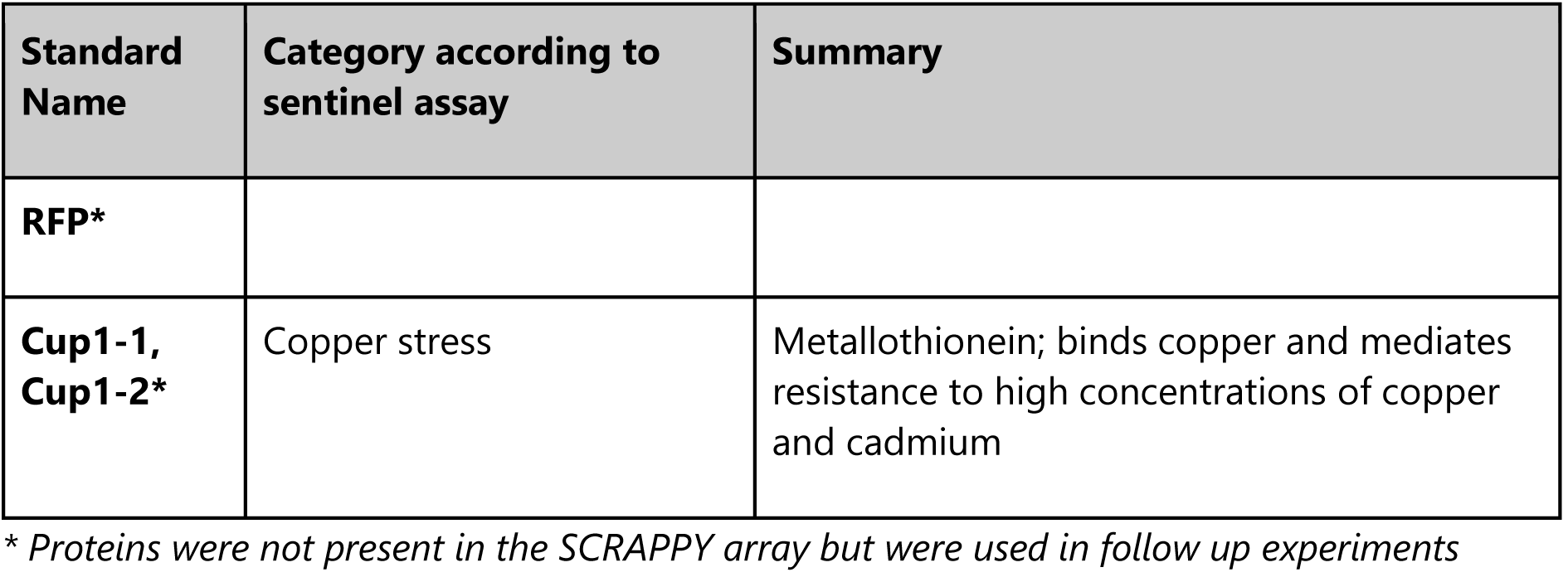
Proteins included into the SCRAPPY array and additional assayed proteins/readouts.

| **Standard Name** | **Category according to sentinel assay** | **Summary** |
| --- | --- | --- |
| **Gcv2** | Amino acid metabolism/catabolism | breaks down glycine by oxidative cleavage |
| **His7** | Amino acid metabolism/catabolism | Involved in histidine biosynthesis |
| **Trp3** | Amino acid metabolism/catabolism | Involved in biosynthesis of aromatic amino acids |
| **Aro2** | Amino acid metabolism/catabolism | Involved in biosynthesis of aromatic amino acids |
| **Lys1** | Amino acid metabolism/catabolism | Involved in lysine biosynthesis |
| **Hom2** | Amino acid metabolism/catabolism | involved in methionine and threonine biosynthesis |
| **Aro3** | Amino acid metabolism/catabolism | Involved in biosynthesis of aromatic amino acids |
| **Car1** | Amino acid metabolism/catabolism | catabolizes arginine to ornithine and urea |
| **Sam4** | Amino acid metabolism/catabolism | Involved in methionine biosynthesis |
| **Sui2** | Amino acid metabolism/catabolism | Upregulated upon translation inhibition |
| **Gcv3** | Amino acid metabolism/catabolism | Glycine catabolism |
| **Rnr4** | DNA damage | Ribonucleotide-diphosphate reductase (RNR) |

| Standard Name | Category according to sentinel assay | Summary |
| --- | --- | --- |
|  |  | small subunit |
| **Pdc1** | Carbohydrate metabolism | Involved in amino acid catabolism and the fermentation of glucose to ethanol |
| **Pyc2** | Carbohydrate metabolism | converts pyruvate to oxaloacetate |
| **Tps3** | Carbohydrate metabolism | involved in trehalose biosynthesis |
| **Tsl1** | Carbohydrate metabolism | involved in trehalose biosynthesis |
| **Pgm2** | Carbohydrate metabolism | catalyzes the conversion from glucose-1-phosphate to glucose-6-phosphate |
| **Ald4** | Carbohydrate metabolism | involved in ethanol metabolism, pyruvate metabolism, acetate biosynthesis and NADPH regeneration |
| **Hxk1** | Carbohydrate metabolism | phosphorylates glucose, mannose, and fructose |
| **Pda1** | Carbohydrate metabolism | involved in acetyl-CoA biosynthesis from pyruvate |
| **Idp1** | Carbohydrate metabolism | catalyzes the oxidation of isocitrate to alpha-ketoglutarate |
| **Tal1** | Carbohydrate metabolism | Non-oxidative pentose phosphate pathway |
| **Hxt3** | Carbohydrate metabolism | Glucose transporter; involved in glucose uptake |
| **Pmi40** | Protein secretion | catalyzes the interconversion of fructose-6-P and mannose-6-P |
| **Hog1** | MAP-kinase | Mitogen-activated protein kinase involved in osmoregulation |
| **Cwp1** | MAP-kinase | Cell wall protein |
| **Mdh2** | Peroxisome | involved in gluconeogenesis and peroxisome matrix protein import |
| **Erg3** | Lipid biosynthesis | C-5 sterol desaturase involved in ergosterol biosynthesis pathway |
| **Erg10** | Lipid biosynthesis | Acetyl-CoA C-acetyltransferase that contributes to ergosterol biosynthesis |
| **Lsc1** | Lipid biosynthesis | Alpha subunit of succinyl-CoA ligase |
| **Pdr5** | Multidrug response | Pleiotropic Drug Resistance, facilitates drug removal from the cell |
| **Dur1** | Nitrogen metabolism | catalyzes degradation of urea to urea-1-carboxylate (allophanate) |
| **Uga1** | Nitrogen metabolism | involved in the 4-aminobutyrate and glutamate degradation pathways; |
| **Gpd2** | Osmotic balance | NAD-dependent glycerol-3-phosphate dehydrogenase |
| **Gpd1** | Osmotic balance | NAD-dependent glycerol-3-phosphate dehydrogenase; key enzyme of glycerol synthesis |
| **Ahp1** | Oxidative stress | Alkyl HydroPeroxide reductase |
| **Grx1** | Oxidative stress | Bifunctional glutathione peroxidase and glutathione transferase |
| **Uth1** | Oxidative stress | Protein involved in mitophagy and cell wall biogenesis |
| **Yhb1** | Oxidative stress | Nitric oxide oxidoreductase |
| **Tsa1** | Oxidative stress | acts as both ribosome-associated and free cytoplasmic antioxidant |
| **Sod1** | Oxidative stress | Superoxide dismutase |
| **Trx2** | Oxidative stress | Cytoplasmic thioredoxin isoenzyme |
| **Ssa1** | Protein folding | HSP70 family ATP-binding protein |
| **Ssa2** | Protein folding | HSP70 family ATP-binding protein |
| **Ssa4** | Protein folding | HSP70 family ATP-binding protein |
| **Ypk1** | Quiescence | Yeast Protein Kinase |
| **Sti1** | Protein folding/quiescence | Hsp70 and Hsp90-binding protein and ATPase inhibitor |
| **Bat2** | Quiescence/amino acid metabolism | Branched-chain amino acid aminotransferase |
| **Tdh1** | Quiescence/Glucose metabolism | Glyceraldehyde-3-phosphate dehydrogenase |
| **Pnc1** | Quiescence | Pyrazinamidase and NiCotinamidase |
| **Hsp12** | Quiescence | Lipid binding protein involved in plasma membrane organization |
| **Hsp104** | Thermotolerance | Adenosine-binding protein disaggregase |
| **Hsp78** | Thermotolerance | Adenosine-binding protein disaggregase (mitochondrial) |
| **Nth1** | Thermotolerance | Trehalase |

| Standard Name | Category according to sentinel assay | Summary |
| --- | --- | --- |
| **Sse1** | Thermotolerance | Peptide and ATP-binding adenylyl-nucleotide exchange factor |
| **Hsc82** | Thermotolerance | Cytoplasmic chaperone of the Hsp90 family |
| **Tps1** | Thermotolerance | Trehalose-6-Phosphate Synthase |
| **Nsr1** | Thermotolerance | Nuclear localization sequence binding protein |
| **Pgk1** | Thermotolerance | Phosphoglycerate kinase involved in glycolysis and gluconeogenesis |
| **Hsp42** | Thermotolerance | Small heat shock protein with chaperone activity; |
| **Aha1** | Thermotolerance | Chaperone-binding ATPase activator involved in protein folding |
| **Hsp26** | Thermotolerance | Small heat shock protein (sHSP) with chaperone activity; |
| **Ydj1** | Thermotolerance | Hsp40-family chaperone |
| **Pre4** | Protein degradation | Beta 7 subunit of the 20S proteasome |
| **Htb2** | DNA/Cell-cycle | Histone |
| **Bmh1** | 14-3-3 protein | Phosphoserine- and DNA replication origin-binding protein |
| **BY4741** | Autofluorescence control | no expression of GFP |
| **Kar2\*** | Unfolded protein response | ATPase involved in protein import into the ER; also acts as a chaperone to mediate protein folding in the ER |
| **Kar2pro-** | Unfolded protein response | Same as above |
| RFP* |  |  |
| Cup1-1, Cup1-2* | Copper stress | Metallothionein; binds copper and mediates resistance to high concentrations of copper and cadmium |

\* Proteins were not present in the SCRAPPY array but were used in follow up experiments

### SOV synthesis

The modified 2’-deoxycytidine derivatives under investigation, N^4^-dodecyl-5-methyl-2,-deoxycytidine (Ala-54) and 3′-amino-N^4^-dodecyl-5-methyl-2′,3′-dideoxycytidine (SOV4), were synthesized by methods developed previously [22,19].

### Thiazolidine synthesis

Mycosidine ((Z)-5-(4-Chlorobenzylidene) thiazolidine-2,4-dione) and 3B-Myc - (4-Chlorobenzylidene)-3-benzoythiazolidine-2,4-dione synthesis was accomplished according to [23].

### Synthesis of ethyl 6-{[[(5-fluoro-2-methylphenyl) amino](imino)methyl]thio-nicotinate 1-oxide hydrochloride (11326083)

Step a) A solution of ethyl 2-chloronicotinate 1 (1.836 g, 1.0 eqv) and urea hydrogen peroxide adduct (UHP) (1.974 g, 2.1 eqv) in methylene chloride (10 mL) was cooled to 0 °C. Then, TFFA (4.15 g, 2.0 eqv) was slowly added to the solution, and the reaction mixture was stirred at room temperature overnight. The reaction mixture was quenched with water and 0.5M water hydrochloric acid solution till pH∼2, and was then extracted by methylene chloride. The organic phases combined were washed by water, saturated NaHCO_3_, dried over MgSO_4_ and evaporated *in vacuo*. The crude ethyl 6-chloronicotinate 1-oxide 2 (1.52 g, 76 %) was recrystallized from ethanol. Mass (EI), *m/z* (*I_relat_*.(%)): 201.6068 [M]^+^ (43). C_8_H_8_ClNO_3_. ^1^H NMR (DMSO-d_6_; 300 MHz; δ, ppm): 8.94 (s, 1H), 7. 81 (dd, 1H, *J* = 8.4), 7.60 (d, 1H, *J* = 8.4), 4.43 (q, 2H, *J* = 7.2), 1.40 (t, 3H, *J* = 7.2).

Step b) To a solution of 5-fluoro-2-methylaniline 3 (0.650 g, 1.0 eqv) in 1.0N water hydrochloric acid solution (6.0 mL) was added ammonium thiocyanate (0.435 g, 1.1 eqv) at 100 °C. The resulting mixture was stirred at 100 °C for 16 h and then cooled to room temperature. The solution was then diluted with cold water and neutralized with 28% ammonium hydroxide solution (pH>7). The precipitate formed was filtered and washed with water and *n*-hexane/diethyl ether to obtain 1-(5-fluoro-2-methylphenyl) thiourea 4 (0.520 g, 46 %) as a white solid. Mass (EI), *m/z* (*I_relat_.*(%)): 184 [M]^+^ (65) C_8_H_9_FN_2_S. ^1^H NMR (DMSO-d_6_; 300 MHz; δ, ppm): 9.24 (s, 1H), 7.26-7.09 (m, 2H), 7.02-6.94 (m, 1H), 2.14 (s, 3H).

Step c) A solution of ethyl 6-chloronicotinate 1-oxide 2 (0.658g, 1.00 eqv) and 1-(5-fluoro-2-methylphenyl) thiourea 4 (0.625 g, 1.03 eqv) in acetone was stirred at room temperature for 12 hours. The precipitate formed was filtered and washed with a small volume of acetone. The crude ethyl 6-{[[(5-fluoro-2-methylphenyl) amino](imino)methyl]thio}nicotinate 1-oxide hydrochloride RCB13083 (0.94g, 83 %) was recrystallized from ethanol. Mass (EI), *m/z* (*I_relat_.*(%)): 349 [M]^+^ (56). C_16_H_17_ClFN_3_O_3_S. ^1^H NMR (DMSO-d_6_; 300 MHz; δ, ppm): 9.11 (s, 1H), 8. 34 (dd, 1H, *J* = 8.4), 8.16 (d, 1H, *J* = 8.4), 7.56-7.46 (m, 2H), 7.18-7.27 (m, 1H), 4.49 (q, 2H, *J* = 7.2), 2.29 (s, 3H), 1.43 (t, 3H, *J* = 7. 2). ^13^C NMR (DMSO-d_6_; δ, ppm): 164.3, 162.4, 156.3, 155.9, 154.5, 140.2, 138.7, 131.8, 131.5, 130.3, 125.6, 125.1, 111.3, 110.6, 109.7, 109.1, 61.1, 17.8, 14.3.

### Creation of the Kar2 promoter-based reporter

The integrative vector pAM978 possessing gene coding for mCherry under control of the *KAR2* promoter was composed of the following DNA fragments : 2237 bp PvuI-EcoRV of pAM784

[24], 115 bp PvuII-BamHI of pGAPZalpha (https://www.thermofisher.com/order/catalog/product/V20520), 504 bp BamHI-NcoI (NcoI was filled-in) of PCR product obtained with primers ScPKAR2U and ScPKAR2L and *S. cerevisiae* DNA as a template, 782 bp PCR product obtained with primers ymCherryU and ymCherryL and pAG426-GAL-ccdb-ymCherry (a gift from Susan Lindquist (Addgene plasmid # 14155; http://n2t.net/addgene:14155; RRID:Addgene_14155)), and 1633 bp SmaI-PvuI of pUC18 *E. coli* vector. This plasmid was integrated into the BY4741 genome. To direct the integration into the *KAR2* locus, the plasmid was digested with XhoI, whose site is unique in the plasmid and is located within the *KAR2* promoter. The following primers were used: ScPKAR2L1 ttCCATGGTATGTTTGATACGCTTTTTC; ScPKAR2U1 aaggatccccatgaactcagca; YmCherryU1 gggttaattaacagtaaaggagaagaagacaacatggcaatc; YmCherryL1 GCAGCCCATCACCACTTTG

### Creation of the *TAT1/2* deletion strain

A double-deletion mutant tat1Δtat2Δ was constructed using tat1Δ strain from Yeast Gene Deletion Collection [9] as a parental strain by disruption of TAT2 gene with *HIS3*. The *TAT2* disruption cassette was obtained via PCR using the primers (TAT2:HIS3-D: TTCATATTTGTTTGTATATACATCTGAGCATTGCGGATCTAAATAGTGTGagcgctaggagtcac TAT2::HIS3-R: AATATTCTACAAAAATAAATTGAACTTGTTTCTTCGGTATTAACACCAGAgcgcctcgttcagaatgac) and pRS313 plasmid as a template. Integration of disruption cassette was assisted by CRISPR Cas9 approach. To do this the plasmid pWS171 [11] possessing the guide RNA assembly cassette and Cas9 encoding gene was used. The duplex of oligonucleotides Tat2Cr-D (GACTGGCTAGGTGAAATCACGGTG) and Tat2Cr-R (AAACCACCGTGATTTCACCTAGCC) were inserted between Esp3I cites of the pWS173 plasmid to direct Cas9 cleavage to the TAT2 locus. The resulting plasmid was used to co-transform tat1Δ strain with the *TAT2* disruption cassette.

## RESULTS

### Selection of proteins for the test-array

Our main goal was to check whether a limited array of *S. cerevisiae* strains producing GFP-tagged proteins would allow observation of clear proteomic signatures for different drugs, as well as to compare them between each other, by monitoring fluorescence of GFP fused to selected sentinel proteins. This selection was primarily based on the paper published by the Picotti group [16], which selected proteins for MS-based identification based on whether a protein was previously observed in MS data, as well as on the literature suggesting whether or not a protein’s abundance characterized a specific cellular process. Our selected panel was smaller than that of the Picotti group, since MS can probe changes in abundance as well as post-translational modifications, and we also had a limitation in method sensitivity due to the inability of the flow cytometer to reliably differentiate low GFP levels from cellular autofluorescence. In short, our panel (Table 2 and Figure 1B) included proteins characterizing amino acid biosynthesis and catabolism, DNA repair, carbohydrate metabolism, lipid biosynthesis, oxidative stress, activation of thermotolerance and quiescence pathways. Several proteins not present in the Picotti group’s final assay were included, due to their high relevance to antifungal drug responses, such as Pdr5-GFP, a 14-3-3 protein involved in regulating phosphorylated protein activity, Bmh1-GFP, as well as Htb2-GFP, a histone which was recently shown by us to allow monitoring of cell cycle perturbations (paper submitted). Using available data on the GFP-fusion collection [13], we only included proteins that were likely to be observable by flow-cytometry.

### Drug selection and treatment optimization

The drugs we selected for testing with the array of strains are listed in Table 2, and included two compounds from the azole class, since these are the antifungals of choice in the clinic and we wanted to compare two compounds with the same target. Another tested compound was SDS, since it is commonly used as a membrane/cell wall stressor and our group has also implicated it in cell cycle perturbations [25]. Bearing in mind that compounds are effective at different concentrations, we used amounts that were based on the minimal inhibitory concentration (MIC). The MICs of the compounds tested were determined for the BY4741 strain (Table 1), which is the parent of the GFP-tagged strain collection used in this study. We assumed that MIC or half of the MIC (MIC/2), at which cells can still proliferate, would be sufficient to induce cellular response. Thus, each drug was tested at two concentrations (MIC and MIC/2), and each test was repeated at least three times.

The time of incubation with the tested compounds should allow the cells to realize their response to the applied stimuli, but should not be too long for technical reasons. Tests with two compounds demonstrated that six hours was an optimal time for the emergence of a clear response (Supplemental materials and Figures S1 and S2). Interestingly, we observed protein leakage from dead cells, as well as slightly increased PI staining of living cells for some compounds, somewhat similar to that observed during laser printing of yeast [26].

### General responses to compounds

A bird’s eye view of the obtained data reveals that a considerable number of the selected proteins show upregulation in response to most of the different tested compounds (Figure 1A). For instance, numerous proteins involved in the heat shock response and proteostasis, as well as proteins involved in carbohydrate metabolism were induced by a number of differing substances. Notably, while most of the tested stressors caused this reaction, which likely relates to the Environmental Stress Response [27], this is not always the case, as demonstrated by azoles (Flu and Vor) and, to some extent, SDS, both of which show almost no or limited induction of chaperones and carbohydrate metabolism genes respectively.

To determine whether the obtained profiles of protein abundance change allowed clear discrimination between compounds of distinct classes, and similarity between compounds of the same class, we using the uniform manifold approximation and projection (UMAP) (Figure 1C) [28] approach to reduce the dimensionality of the data. This approach, somewhat similar to principal component analysis, allows visualization of multidimensional data on a two-dimensional representation. We observed that different concentrations of the same drug are usually close together on the UMAP, as well as drugs from the same class.

### Specific proteomic response of tested drugs Azoles

An important question was whether the assay revealed specific cellular responses to drugs with known mechanisms of action. Both fluconazole and voriconazole caused robust upregulation of Erg3-GFP and Erg10-GFP, unlike all of the other tested drugs (Figure 1D, Figure 2A, B). This was in good accordance with the known mechanism of action of azoles, which involves inhibition of Erg11, and subsequent changes to the lipid metabolism. Importantly, deletion of *ERG3* is known to cause resistance to azoles [29], which we confirm independently (Figure S3). It is thought that this resistance emerges via preventing formation of toxic intermediate products of ergosterol biosynthesis, i.e. in this case increased amounts of Erg3 seems to exemplify a maladaptive cellular response.

**Figure 2.**
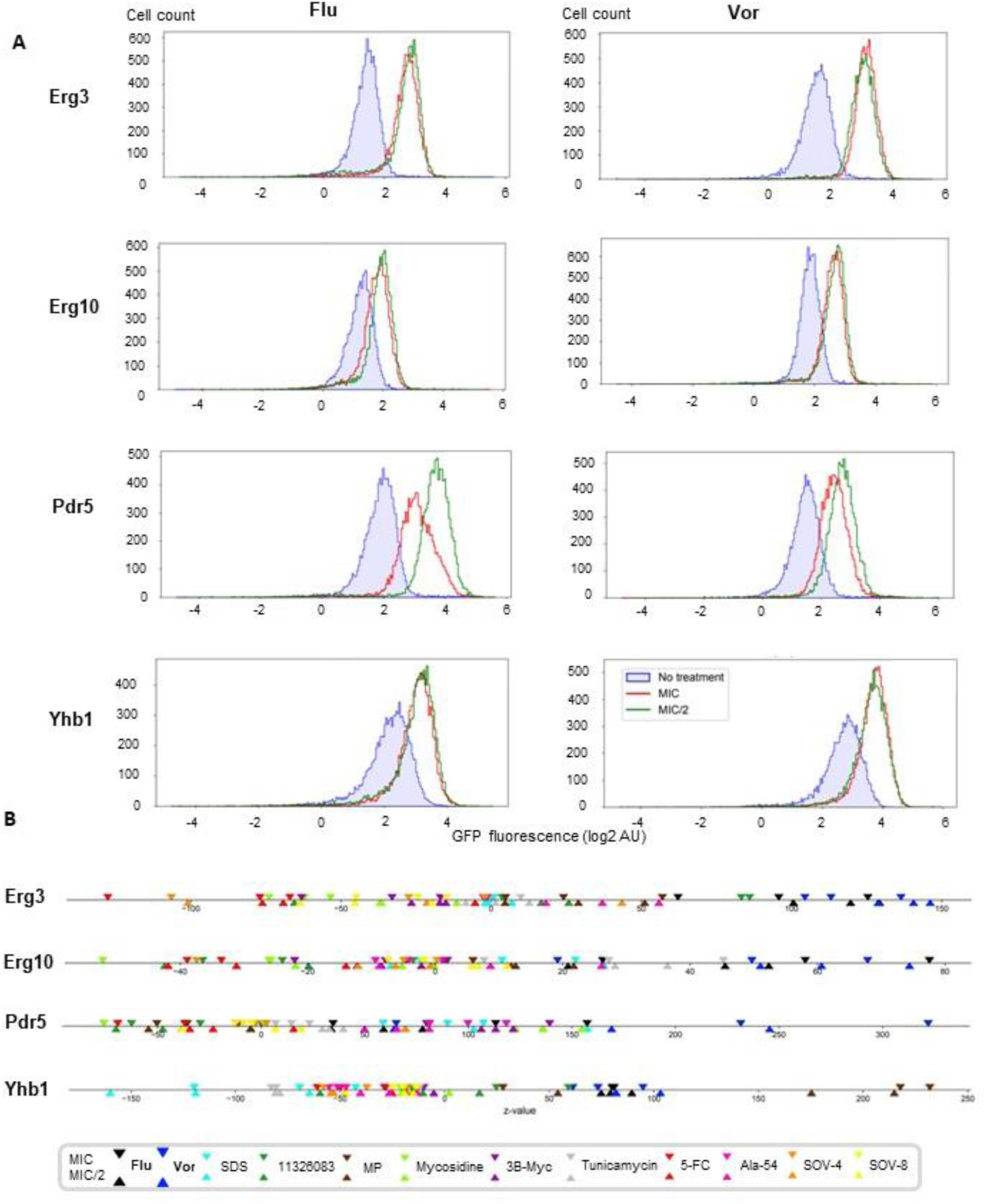
SCRAPPY demonstrates the ability of azoles to upregulate lipid biosynthesis enzymes and drug transporters. (A) Cytometric data on the changes in protein level; (B) Comparative unidimentional plot of z-score changes for all tested compounds

As expected, azoles also demonstrate induction of the drug transporter Pdr5 (Figure 2A,B), which is known to modulate their toxicity by facilitating drug efflux [30] (Figure S3), although this response is not azole specific and can be observed for several other compounds in our dataset. We also observed induction of the low-affinity glucose transporter Hxt3 (Figure 1C), which is induced by both azoles and thiazolidines (see further), however deletion of *HXT3* has no effect on azole toxicity (Figure S3B).

Interestingly, in line with the numerous reports of oxidative stress during azole treatment of fungi [31–36], we observed clear responses for Yhb1 and (to a lesser extent), Tsa1 (Figure 1C, Figure 2A, B).

### Divalent metal ion-transporting molecules – classic and novel

Another substance that we tested was a component of the commonly used anti-dandruff drug zinc pyrithione, 2-mercaptopyridine-*N*-oxide (MP), also known as pyrithione. Zinc pyrithione is known to cause copper and zinc influx into fungal cells [2, 37]. This substance, as well as another chemical, synthesized in the course of this project (termed 11326083), is related to a recently tested anti-mycobacterial compound shown to induce copper influx [38]. Both MP and 11326083 demonstrated a very similar response, suggesting that they share a mechanism of action, although several proteins induced by MP were induced less efficiently by 11326083. As expected during copper influx [39], we observed a considerable oxidative stress response (increased levels of Ahp1, Sod1, Trx2) (Figure 3A,B), as well as the strongest observed induction of the chaperone Ssa4, suggesting proteotoxic stress, likely due to protein oxidation. Notably, both of the compounds seem to cause downregulation of Lys1 (Figure 1D,E), which is unique among the presented dataset. Notably, this is in line with the known ability of pyrithione to inactivate Fe-S proteins via copper influx [38], including aconitases, which are involved in the early stages of lysine biosynthesis [40].

**Figure 3.**
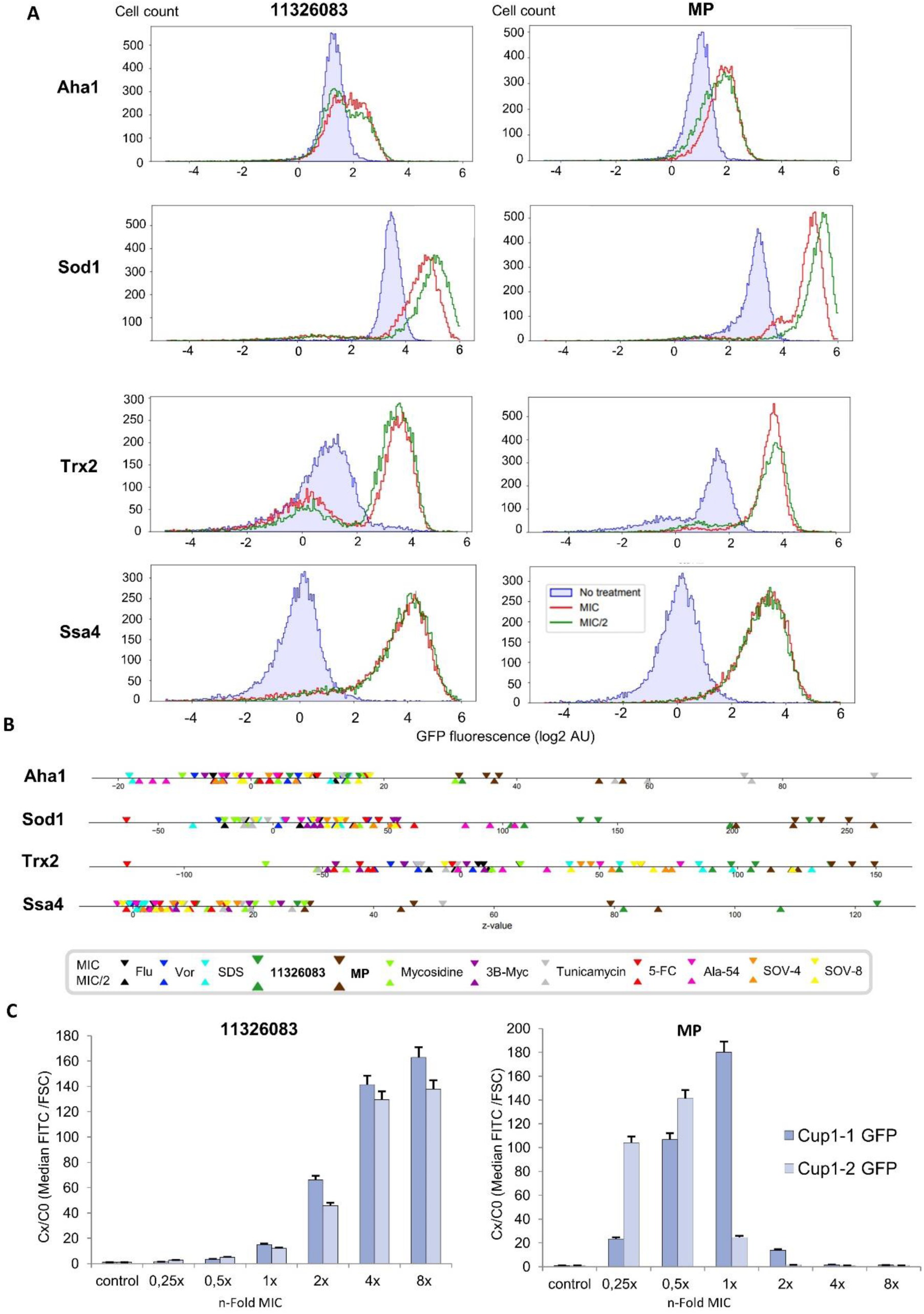
MP and 11326083 demonstrate strong induction of the oxidative stress and proteotoxic response, as well as specific induction of copper detoxifying enzymes. (A) Cytometric data on the changes in protein level ; (B) Comparative unidimentional plot of z-score changes for all tested compounds; (C) Change in the level of Cup1-GFP proteins in response to compound treatment (MICs for 11326083 and MP are noted in Table 1).

To confirm that copper influx was the likely mechanism for 11326083, we tested for the induction of Cup1-1-GFP and Cup1-2-GFP, which are metallothioneins involved in copper detoxification. These proteins showed robust induction in the presence of 11326083 and MP (Fig. 3C). Notably, we could observe that protective enzyme induction is much weaker for MIC-equivalent concentrations of 11326083 and the reaction persists to much higher MIC-equivalent concentrations. This suggests that despite using MIC-equivalent concentrations, the effects of 11326083 are milder than that of MP. These results and the differences between MIC-equivalent profiles of 11326083 and MP suggest the possibility that while the core mode of action of two compounds likely involves copper ion influx, they probably differ in some more subtle way.

### SDS

SDS sensitivity has been extensively used as a stressor, which reveals impairment of the cell wall and/or plasma membrane [41–43] and it has also been implicated in control of the cell cycle [25, 44]. In accordance with the previously reported detection of reactive oxygen species using dihydroethidium staining [44], we observed induction of Trx2 (Figure 1D,E, Figure 3B). However, despite the fact that SDS-sensitivity is dependent on aromatic amino acid biosynthesis [42, 44], cells did not upregulate the corresponding proteins, even though alkylated cytidines did so (see further).

Interestingly, SDS was not highly effective in causing upregulation of the chaperones and carbohydrate metabolism proteins, which is observed for most other tested compounds, apart from azoles.

### Thiazolidines

For comparison to other drugs, we also performed profiling for two thiazolidine derivatives, which were recently reported in [23]. While thiazolidines did not provide as distinct a profile as that of azoles, SDS or copper-influx compounds in terms of uniqueness, mycosidine and 3B-Myc showed induction of the glucose transporter Hxt3 (only at low drug concentration for mycosidine) and drug transporter Pdr5 (Figure 1D,F and Figure 4), which is somewhat similar to azoles (Figure 1C, Figure 4B). Interestingly, Hxt3 deletion increased mycosidine activity [23].

**Figure 4.**
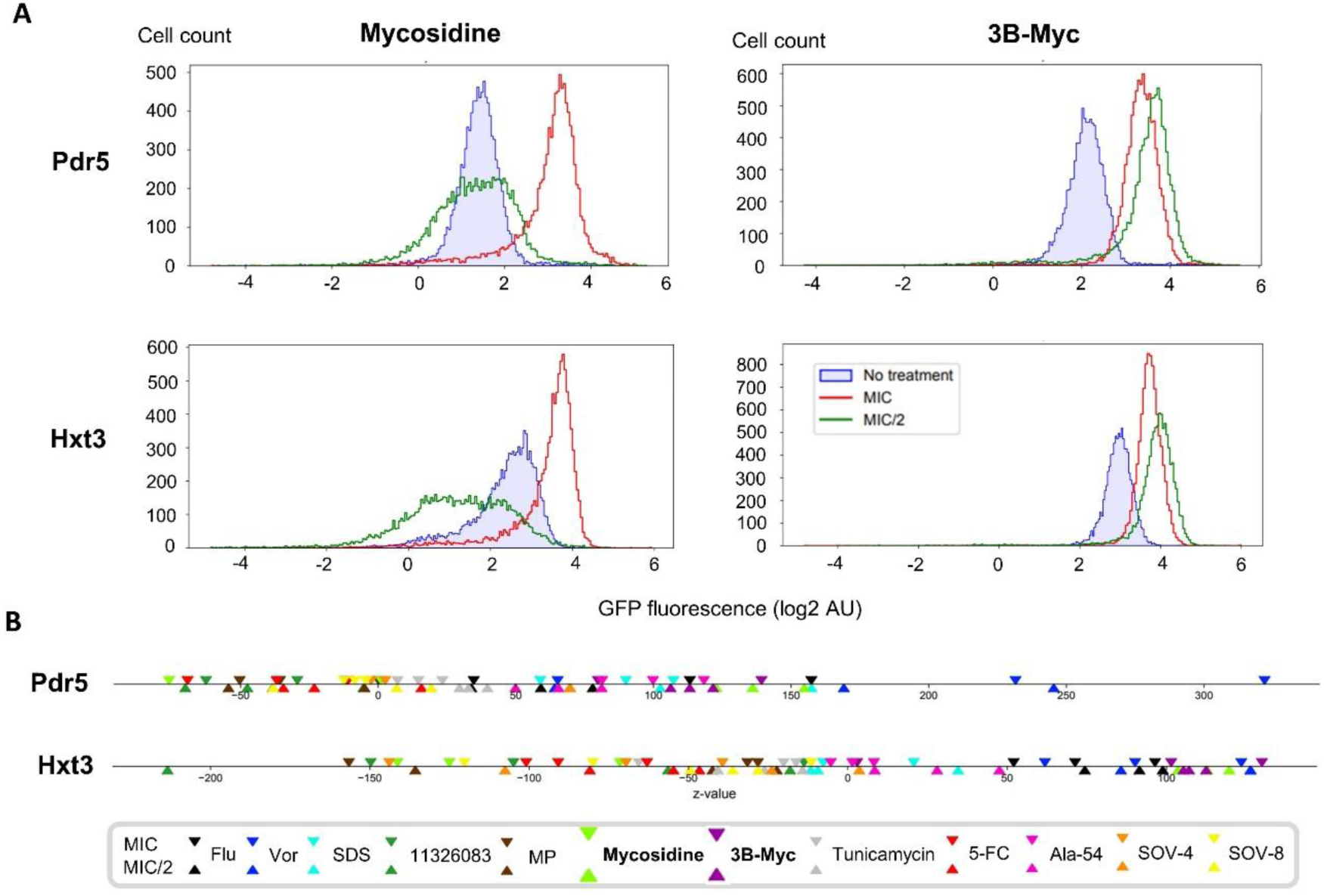
Mycosidine and its related compound 3B-Myc specifically induce Hxt3 and Pdr5 levels, showing similarity to azoles. (A) Cytometric data on the changes in protein level; (B) Comparative unidimentional plot of z-score changes for all tested compounds.

Importantly, both Pdr5 and Hxt3 are also induced by azoles (Figure 1D), with Pdr5 having an important role in azole toxicity (Figure S3), which is not the case for mycosidine [23]. As noted above, *HXT3* deletion did not affect toxicity of Vor (Figure S3), but influenced resistance to mycosidine, which suggests that for thiazolidines, Hxt3 plays an important role in the mode of action. Interestingly, it has previously been reported that thiazolidine derivatives are efficient blockers of glucose transport [45], though, notably, this paper was later retracted. However, several more reports implicate thiazolidine-based molecules in glucose transport [46, 47]. Interestingly, the thiazolidine derivatives used in the clinic are considered to be peroxisome proliferator-activated receptor (PPARγ) agonists, while these receptors are absent in yeast. This suggests that the effects of thiazolidines may be mediated by other targets in yeast and possibly mammals.

### Tunicamycin

Tunicamycin is a known ER-stressor, which inhibits Alg7, a uridine-diphosphate-N-acetyl-glucosamine-1-P transferase catalyzing an essential step of synthesis of carbohydrate chains, which are attached to asparagine residues of the nascent polypeptide chains translocated into the ER lumen. This is crucially required for folding the majority of proteins in this compartment. As could be expected, we observed specific induction of Pmi40 (Figure 5A,B), which converts fructose-6-phosphate into mannose-6-phosphate required for protein glycosylation in the secretory pathway, and Pre4, which is a subunit of the proteasome, involved in the degradation of damaged ER proteins. Interestingly, tunicamycin also causes strong induction of Tsa1 (Figure 5), a peroxiredoxin and a partner of Hsp70 which facilitates interaction with oxidatively damaged proteins [48]. Tsa1 was also induced, albeit less dramatically, by the copper-transporting compound MP. We also detected specific induction of Aha1 (Figure 5) and Ydj1 (Figure 1D), which are partners of Hsp90s and Hsp70 respectively.

**Figure 5.**
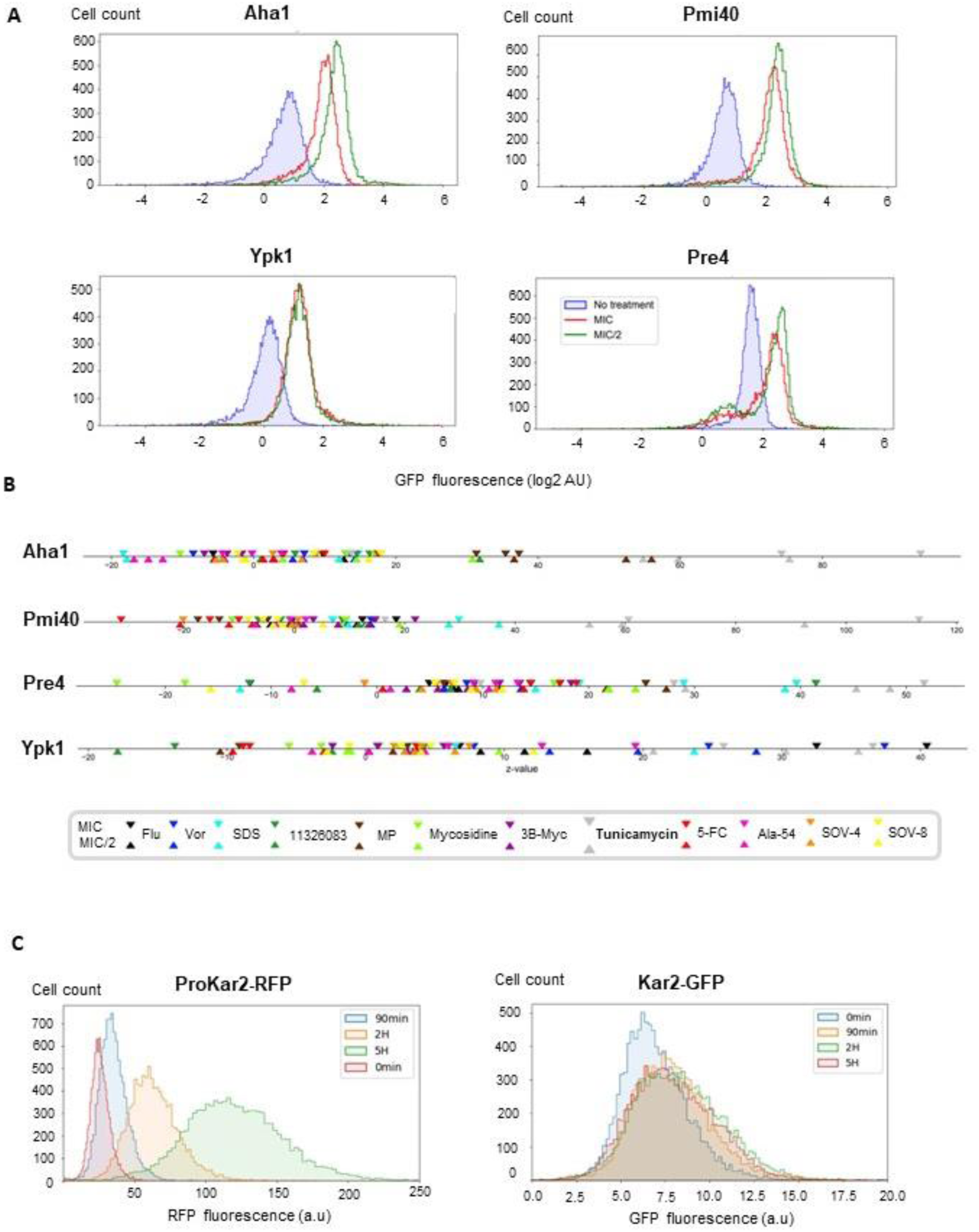
Tunicamycin induces specific increases of Pmi40 and Pre4 and induces KAR2 expression, which is detected more reliably using a promotor-activity based sentinel, rather than a mislocalized GFP-fusion protein. (A) Cytometric data on the changes in protein level. (B) Comparative unidimensional plot of z-score changes for all tested compounds; (C) Responses of KAR2-promoter reporter and Kar2-GFP fusion protein to tunicamycin treatment. Cells harboring RFP under control of the KAR2 promoter (right) or containing the Kar2-GFP fusion protein were grown in medium containing tunicamycin at MIC concentration (2 µg/ml). While the MIC of the Kar2-GFP strain was 2-fold lower than that of the KAR2-promoter strain, the response of Kar2-GFP was strongest at the presented concentration.

These observations highlight the distinct type of proteotoxic oxidative damage triggered by tunicamycin. Tunicamycin also provides a good test case for determining optimal setups and options for future SCRAPPY modifications. While using the available GFP-fusion collection is highly convenient, some proteins are ill suited for assay as a C-terminal fusion. For instance, these are proteins possessing C-terminal localization signals, e.g., ER retention signals HDEL or KDEL. One such protein of interest, which might be used as a “sentinel” of the unfolded protein response [49] is Kar2, which is an ER chaperone. Notably, its GFP-fusion (or at least the one present in the available GFP-fusion collection) is not localized to the ER, as should be the case for a fully functional protein. This does not, at least in theory, prevent its use as a promoter activity readout.

So, we decided to compare the behavior of this protein with an alternative sentinel setup, where a fluorescent protein (mCherry) was placed under the control of the Kar2 promoter (*P_KAR2_*). Of course, this changes the sentinel readout to be one of transcriptional activation, rather than of increased protein abundance. After treatment with tunicamycin, we observed that while Kar2-GFP did show a weak increase, the response of *P_KAR2_-mCherry* was far stronger (Figure 5C).

This demonstrates the usability of promoter-based sentinels in the SCRAPPY-like approach, and also that some sentinels can benefit from using promoter-based reporter constructs as opposed to fusion proteins.

### Non-homogeneous change in mitochondrial protein levels

Up to now, we did not utilize the ability of flow cytometry to provide single-cell level data. Notably, this method allows addressing whether different populations of cells in a suspension can respond differently to a treatment. To assay, whether this was common, we analyzed fluorescence distributions for each protein and screened them manually in order to detect cases where the distribution of cells was unimodal in the control, and markedly non-normally distributed and preferably, multimodal, after treatment. While most proteins responded in a homogenous manner, we did observe a somewhat bimodal or widely distributed response for the mitochondrial protein Ald4-GFP. In response to a range of treatments (5-FC, copper-transporting drugs and thiazolidines), some of the cells retained their baseline level fluorescence, while a considerable portion of cells exhibited increased levels of the protein. Idp1 which is also a mitochondrial protein, exhibited a similar response, albeit only after treatment with thiazolidines (Figure 6). In order to elaborate on this phenomenon, we used fluorescent microscopy to observe the localization of the respective proteins upon treatment with mycosidine. Under control conditions, the mitochondria were single long string-like structures, while after treatment, for both proteins, we observed obvious change of mitochondrial morphology, as well as heterogenous changes of protein level. This fully supports the data obtained by flow cytometry.

**Figure 6.**
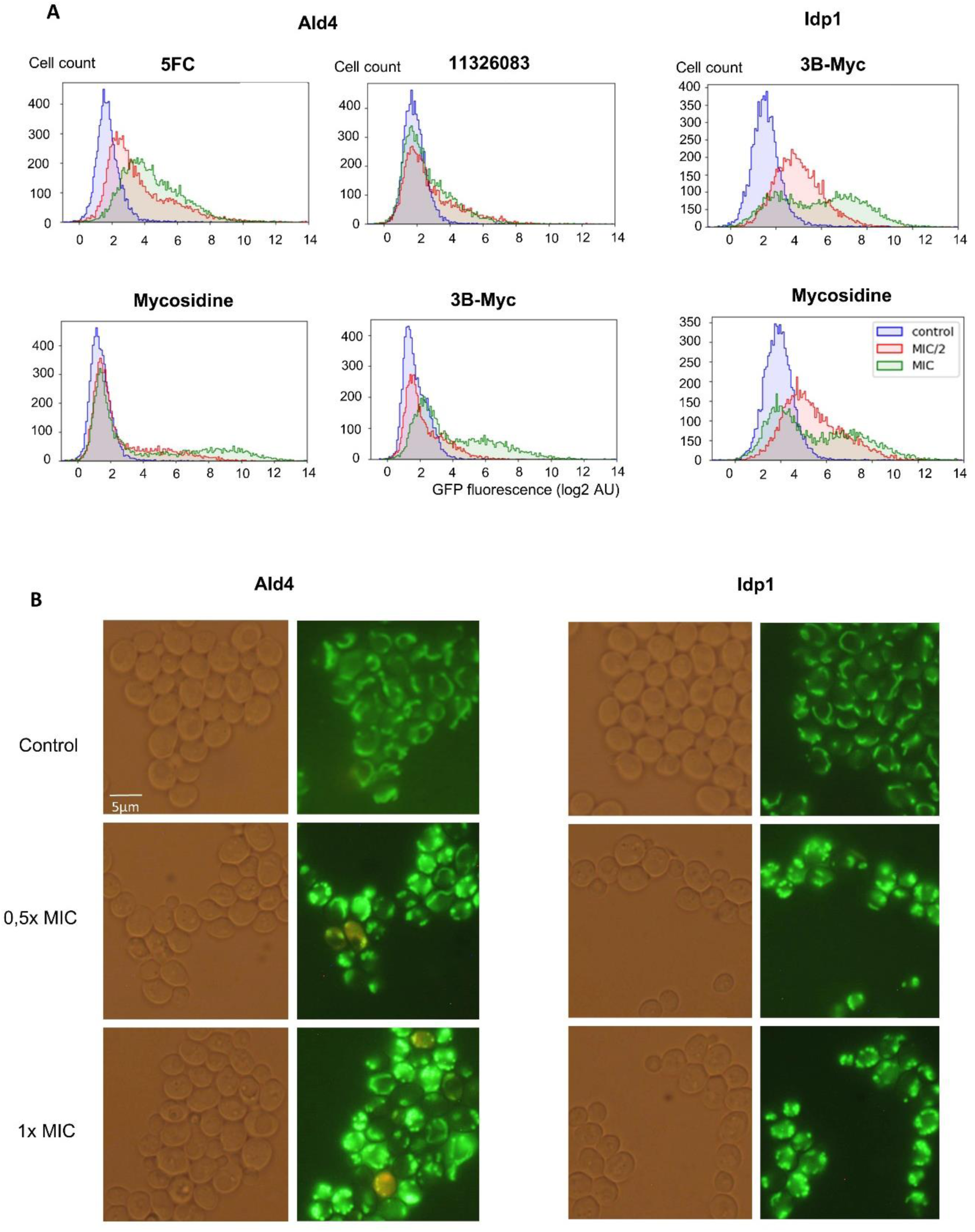
Mitochondrial proteins Ald4 and Idp1 display non-homogenous changes in response to treatment. (A) Cytometric data on the changes in protein level; (B) Fluorescent microscopy of the same cells used for the cytometric analysis under visible light and GFP fluorescence conditions.

### Monitoring effects on the cell cycle

One of common uses of cytometry is to measure the DNA content of individual cells to characterize the cell cycle. Our recently published method on the use of GFP-tagged histones for cell cycle characterization [50], as well as the inclusion of Htb2-GFP into the SCRAPPY panel, allowed us to easily identify compounds which had an effect on the cell cycle. Due to the use of propidium iodide staining, we could also observe only living (PI negative cells). As expected, though all of the tested compounds stop cell division, the way in which the compounds do this is not identical. Usually, actively dividing cells exhibit a smaller population of cells with a single complement of DNA (1C) and larger population of cells with replicated DNA (2C), as well as some cells that are replicating their DNA (S). We observe that azoles do not change the distribution of histone abundance (Fig. 7A), while thiazolidines and copper ionophore compounds both caused cellular arrest with histone amounts corresponding to 1C. To our knowledge, this is the first report of such an activity for copper ionophores. As expected, and reported previously, 5-FC and tunicamycin caused accumulation of cells in 2C population[50] and [51]. SDS seems to cause a delay in the S-phase at lower concentrations and causes cell to pause at 2C if used at MIC. SOV4 resulted in most living cells exhibiting histone abundance corresponding to 1C. Interestingly, since both SDS and SOV4 caused considerable cell death, we could observe that inclusion of PI-positive cells revealed a sub 1C peak in the distribution, indicating histone leakage (Fig. 7B). Interestingly, for other compounds where a sub-1C peak was observed, removal of dead cells from the analysis did not remove the peak. Thus, copper ionophores and Myc can cause reduction of histone amounts, which might be due to DNA degradation in cells that are impermeable to propidium iodide. This is often interpreted as a sign of apoptosis-like death [31, 52].

**Figure 7.**
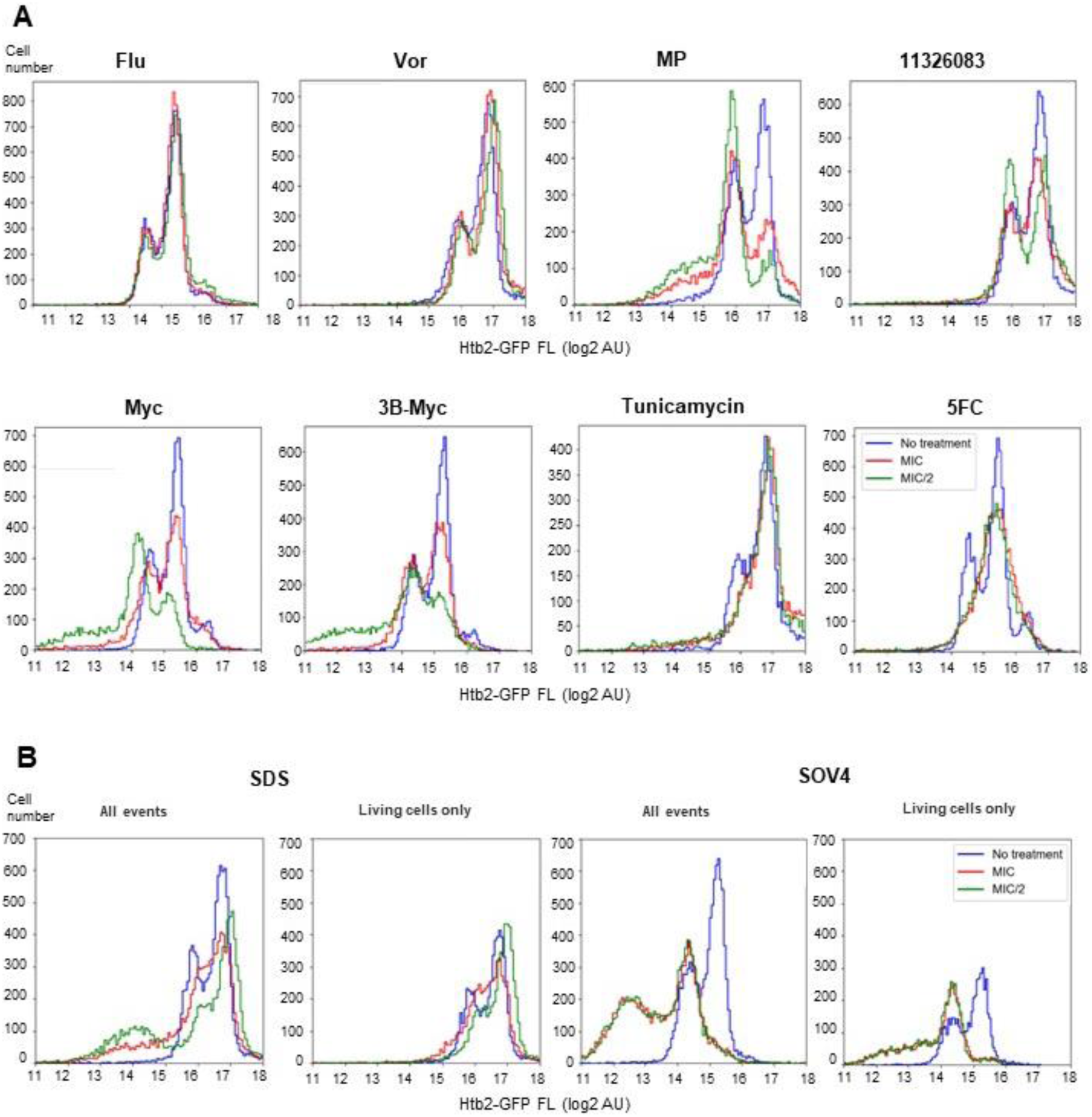
Histone abundance quantification using Htb2-GFP reveals effects of antifungal compounds on the cell cycle state of individual yeast cells. (A) Cytometric data on the changes in Htb2-GFP fluorescence in PI-negative cells; (B) cytometric data on the changes in Htb2-GFP fluorescence in PI-negative cells (living cells only panels) and all the cells (All events panels).

### SCRAPPY provides key information for the characterization of the unknown mode of action of N4-alkyl-cytidines

To demonstrate the usefulness of the SCRAPPY profiles for deeper exploration of a specific class of chemicals with an unknown mode of action, we used it to profile the effects of a set of N4-alkyl-cytidines (Ala-54 and SOVs) which can be used to protect works of art from the degradation caused by fungal pathogens[19, 22]. As a reference, we also obtained SCRAPPY profiles for another nucleic-acid-based antifungal that is used in the clinic, 5-FC. We tested 3 SOV compounds, which had differing MIC’s, including one that had hardly any fungitoxic activity compared to the others, but was used at near the same concentrations for comparison (Table 1). The SCRAPPY profiles are presented, and when these are compared with the other compounds, one can observe distinct features: (1) there is notable upregulation of some proteins in the amino acid biosynthesis cluster, namely Hom2, Aro3, and somewhat less vividly, Trp3 (2) Upregulation of Trx2 (Figure 8A).

**Figure 8.**
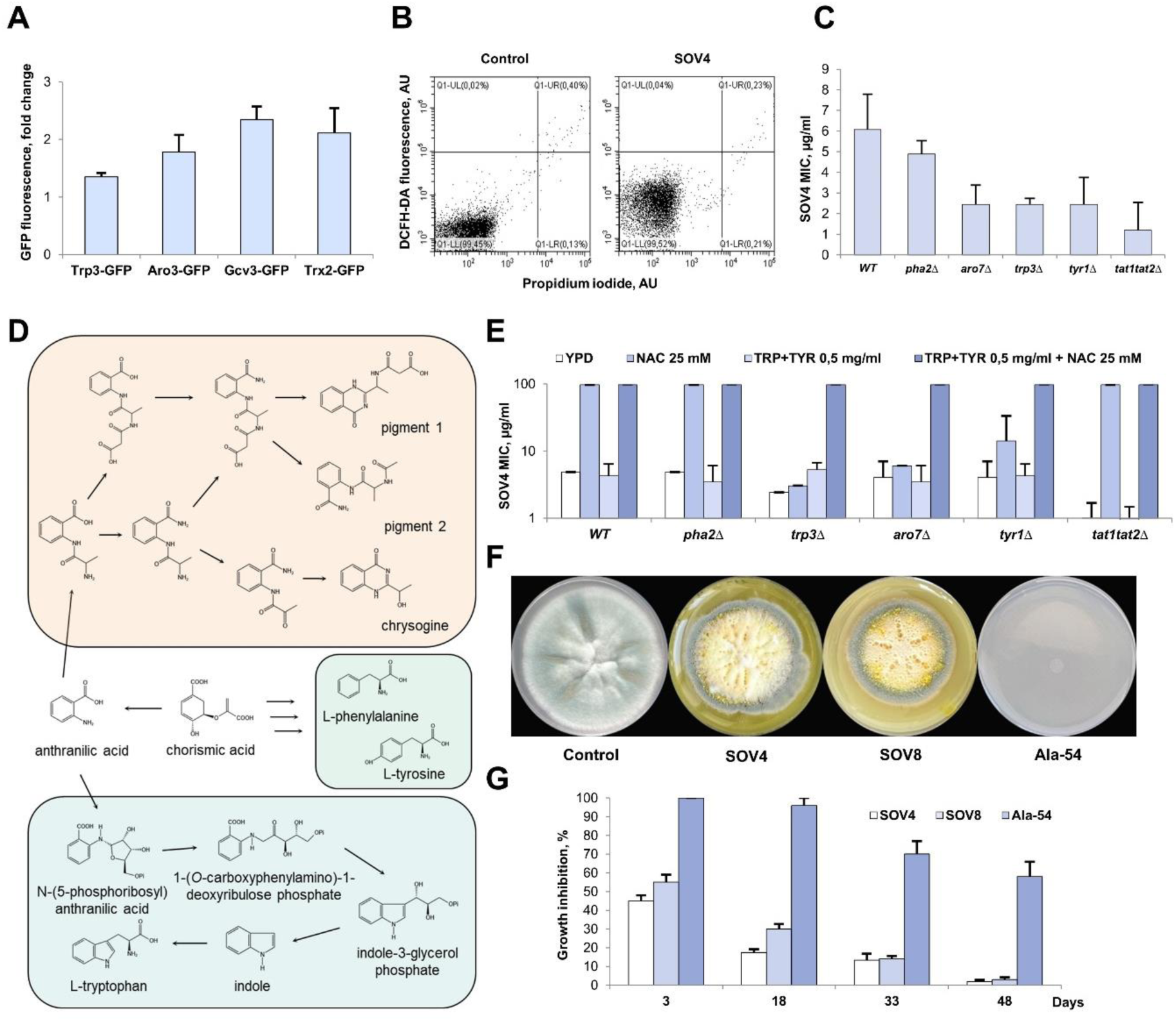
N4-alkyl-cytidines induce production of proteins related to aromatic amino acid biosynthesis and oxidative stress, both of which processes are involved in their toxicity. (A) Cytometric data on the changes in protein level for SOV-4 taking into account wild-type cell autofluorescence; (B) Cytometric data for DCFH-DA staining. SOV4 raises ROS level in 15 minutes after addition to PI-negative cells. (C) Yeast strains with deleted genes involved in aromatic amino acid biosynthesis or transport display increased sensitivity to SOV-4 (D), which can be effectively mitigated by addition of antioxidants only in the wild type or aromatic amino acid transport mutant.

SOV-4 and Ala-54 both also resulted in upregulation of Pdr5. Notably, while Pdr5 has no effect on SOV-4 activity, Ala-54 becomes nearly as toxic as the other derivatives only after Pdr5 deletion (Figure S4A). The observation that the role of Pdr5 is influenced by quite a small change (exchange of a -NH2 for an -OH in the sugar moiety) in the chemical structure is interesting and somewhat unusual.

In order to determine, which specific impairments of the amino acid biosynthesis machinery could affect SOV activity, we tested a set of strains with deletions in the amino acid biosynthesis pathways (Figure S4B) and observed that it was aromatic amino acid biosynthesis that caused the strongest effects, leading to increased sensitivity (Fig. 8C). Importantly, transport of aromatic amino acids also played a role in the toxicity of SOVs, as deletion of the Tat1 or both Tat1/2 permeases increased sensitivity (Figure 8C). Interestingly, addition of Trp and Tyr on its own decreased sensitivity to SOV only in the *trp3*Δ strain (Figure 8D). Because our observations showed induction of Trx2, a component of the oxidative stress response, we used two additional methods to confirm the presence of oxidative stress, and its causative role in SOV toxicity. Firstly, we demonstrated that short term treatment with SOV increased staining with the redox-sensitive dye DCFH-DA (Figure 8B). Secondly, we assayed the role of antioxidants on SOV toxicity. Sensitivity of the wild-type strain, as well as the sensitive mutant strains could be reduced by addition of antioxidants N-acetyl cysteine (NAC) (Figure 8D), as well as ascorbic acid and glutathione (Figure S4C).

Most interestingly, NAC alone was unable to mitigate the effects of SOV in aromatic amino acid biosynthesis mutants, except for *pha2Δ*. However, NAC was highly effective in reducing SOV activity in the aromatic amino acid transport mutants. This suggests that mitigation of SOV-mediated damage requires both an antioxidant and internal synthesis of Tyr/Trp (Figure 8D).

Because previous work had shown that various fungi infecting works of art are sensitive to SOV, we decided to examine whether there are indications that in these fungi, the aromatic amino acid pathway was also influenced by SOV. For this we used the filamentous fungus *Penicillium chrysogenum*, strain STG-117, which was isolated from an icon from the Russian State Tretyakov Gallery [53]. The growth of this strain was previously shown to be completely inhibited by growth on solid CDA medium with 0.7 mM SOV4, or SOV8, or Ala-54 [19]. In order to observe effects of growth on subinhibitory concentrations of the compounds, we used a lower concentration (0.2 mM). While this concentration was completely inhibitory for Ala-54, highlighting the differential sensitivity to SOVs between *Penicillium* and *Saccharomyces* (Fig. 8), slowed growth on the other SOV derivatives (that were also effective in *S. cerevisiae*) was accompanied by accumulation of yellow droplets on the mycelium surface, which are known to contain the pigments chrysogine and others in the same pathway (Fig. 8) [54].Yellow staining was observed both in the mycelium itself, in the exudate (droplets released on the surface of the mycelium) and also penetrated into the agar medium. Importantly, chrysogine and the other yellow pigments of its biosynthetic pathway are synthesized from anthranilate, a precursor metabolite in the tryptophan biosynthesis pathway, and a derivative of chorismate required for the biosynthesis of tyrosine and phenylalanine (Fig. 8). Notably, chrysogine has also been demonstrated to exhibit antioxidant capacity [55]. The obtained data indicates that the effects of SOV on the aromatic amino acid synthesis pathway are conserved among different groups of fungi, and possibly suggests that accumulation of antioxidant metabolites can act as a mechanism of SOV resistance in *Penicillium*.

These results are highly concordant with the reported, yet mostly underappreciated, cytoprotective antioxidant function of tyrosine and tryptophan residues in transmembrane proteins [56] and are highly likely to be relevant for a wide range of phenotypes attributed to deletions in the aromatic amino acid biosynthesis pathway[57].

## Discussion

This study is the first test of an approach to limited chemo-proteomic profiling that provides information on single cells. The assay an panel we have tested shows promise as a methodology for rapid testing of various bioactive substances and treatments in terms of understanding their cellular response, and the array of sentinel proteins can be improved and/or tailored to more specific tasks. We show that the profiles obtained by SCRAPPY are reproducible and exhibit high similarity between drugs that are known to or are supposed to have similar mechanisms of action (Figure 1C). Extensive use of SCRAPPY for chemo-proteomic profiling can quite easily provide a database that would allow placing novel substances on a landscape of profiles obtained for previously tested treatments. It is also useful for pinpointing a novel substance with a distinct and interesting profile, which would warrant deeper characterization, as exemplified by SOV compounds in our study (Fig. 8).

While the mechanism of action for SOVs is still far from being completely clear, we have conclusively demonstrated that these compounds cause a specific type of oxidative stress (via a currently unknown route), which differs from other oxidative-stress inducing compounds and that aromatic amino acids are required to survive this stress even in the presence of antioxidants. Further study is required to explore how alkylated cytidines trigger oxidative stress and why it specifically affects aromatic amino acids, but a likely explanation is that the produced type of oxidative species localize in the membrane, where aromatic amino acids are one of the more vulnerable entities.

One clear challenge for the SCRAPPY method that is currently unsolved is the detection of proteins with low expression levels. This can be approached by testing new brighter fluorescent proteins, as well as those with spectra in which the background fluorescence is weaker. Another important issue is the time of incubation with the drug, because as of now, we used a six hour incubation, which can, potentially, miss transient and rapid cellular responses, such as those reported for calcium levels in yeast cells [58].

The notable advantages of SCRAPPY are the speed and ease of gathering and analyzing the data, low cost of the assay and moderate consumption of novel bioactive substances, which are often available in limited quantities during the development of lead compounds. These features allow testing multiple treatment conditions for a single substance, which is sometimes prohibitively expensive for omics-based methods. The ability to measure the amount of protein in single cells provides the opportunity of monitoring the effect of compounds on the cell cycle (Figure 7). It also allows monitoring divergent responses in a cell population, which were demonstrated in this paper for mitochondrial proteins. This allows detecting cell-to-cell heterogeneity, which might be important for identifying populations of drug resistant cells, or persistors [5], as well as subtle responses which would otherwise remain unnoticed due to data averaging.

In conclusion, the SCRAPPY method holds significant promise as a rapid and cost-effective tool for chemo-proteomic profiling at the single-cell level, offering the potential to advance our understanding of cellular responses to bioactive substances. Its ability to uncover subtle responses and identify divergent cell populations makes it a valuable and novel asset in the field of drug discovery and cellular biology.

## SUPPLEMENTARY DATA

### Optimization of SCRAPPY conditions

To allow cells to realize their response to the applied stimuli, we used an incubation time of 6 hours. We arrived at this time by testing two compounds for two to seven hour time periods. We observed that while the response to one of compounds developed in 4 hours and then remained relatively unchanged, the other compound showed a clear result only after 6 hours (Figure S1). Incubating for longer than 6 hours would be technically incovenient, while shorter time might not be enough to achieve a clear reaction from cells that were under stress and needed to change their protein expression pattern. We acknowledge that some responses might be achieved more rapidly and could be gone by the time of the assay point.

One of the important benefits of flow cytometry is that it allows easy detection and subsequent gating of dead cells according to their staining by propidium iodide. This stain, in most cases, cannot enter cells with an intact membrane, thus the data we present were only calculated for live cells, since dead cells exhibit leakage of numerous intracellular proteins (Figure S2A).

Important observations were made when using PI for this purpose. For most compounds, the majority of cells did not exhibit increased PI staining, while a fraction exhibited strong staining (Figure S2B), indicating that the cells are dead and permeabilized. For most compounds included in this analysis, the number of dead cells was less than 5%, while tunicamycin had ∼30% dead cells and alkylated cytidines and SDS - 30-70% of the cells (Figure S2C), which still did not prevent us from obtaining data on the cellular response.

Notably, in several cases we observed emergence of weak PI staining (Figure S2D), somewhat similar to that observed during laser printing of yeast [26]. This suggests that among the tested compounds, MP, and, to a lesser extent, 11326083, Flu and Vor caused a minor increase in the permeability of cells to PI. Notably, because the vast majority of cells exhibit this effect, and, based on the literature and our own experiments, azoles and 11326083 are not fungicidal, we can assume that such weakly PI-positive cells are alive. This suggests changes to the overall permeability of cells, which could be leveraged for use in combination drug therapy and might explain some of the reported synergies of azoles with other compounds.

### List of direct links to deposited cytometry data

http://flowrepository.org/experiments/6895

http://flowrepository.org/experiments/6888

http://flowrepository.org/experiments/6887

http://flowrepository.org/experiments/6889

http://flowrepository.org/experiments/6890

http://flowrepository.org/experiments/6891

http://flowrepository.org/experiments/6893

http://flowrepository.org/experiments/6892

http://flowrepository.org/experiments/6896

http://flowrepository.org/experiments/6898

http://flowrepository.org/experiments/6897

http://flowrepository.org/experiments/6895

## AUTHOR CONTRIBUTIONS

EG performed SCRAPPY experiments with all the listed compounds, created figures and edited the manuscript, VAB performed studies of SOV after initial SCRAPPY data collection, FR analyzed the data and created analysis scripts, created figures, OVM created constructs and obtained the data on induction of Kar2 promoter and protein, OBR and APE synthesized the compound 11326083 and performed preliminary testing of its activity, YMS edited the manuscript and analyzed the data, IBL synthesized the 3B-Myc and mycosidine compounds and analyzed the data, LAA, MVJ and DAM synthesized SOVs and analyzed the data, VVK devised and implemented the successful strategy in the deletion of aromatic amino acid transporter gene, MOA created the constructs for the Kar2 promoter reporter, took part in conceiving the project and contributed funding, AIA conceived the project, drafted and edited the manuscript, contributed funding.

## ACKNOWLEDGEMENTS

Flow cytometry was performed at the Shared-Access Equipment Centre “Industrial Biotechnology” of FRC “Fundamentals of Biotechnology” (RAS)

