## supplemental figures for "SCRAPPY - a single cell rapid assay of proteome perturbation in yeast uncovers a joint role of aromatic amino acids and oxidative stress in the toxicity of lipophilic nucleoside analogs": Supplemental figures.pdf

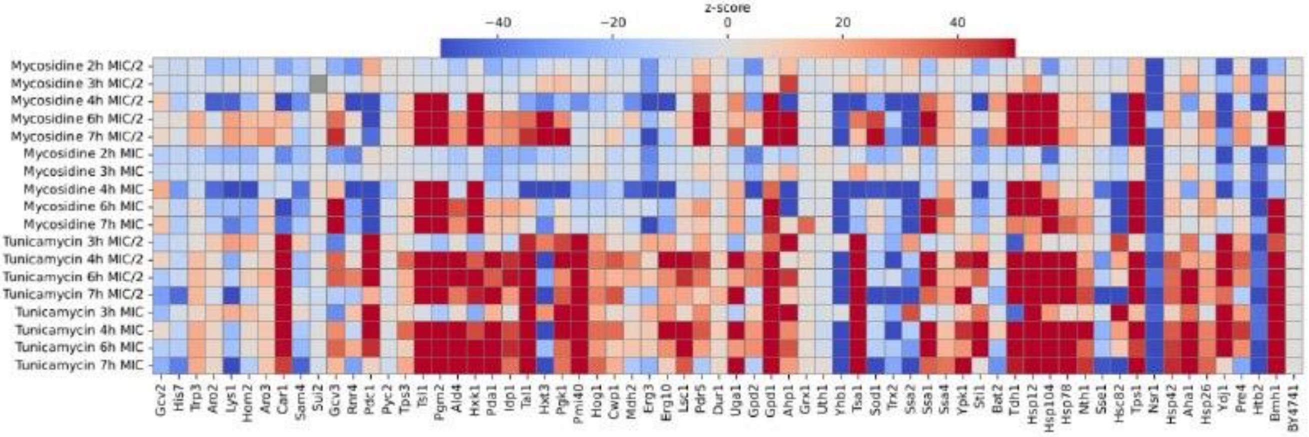


**Figure S1. Optimization of incubation time for mycosidine and tunicamycin.** Cell of the SCRAPPY array were incubated with compounds for the noted amount of hours (h) at the noted concentration and then subjected to flow cytometry as noted in the Materials and Methods section.

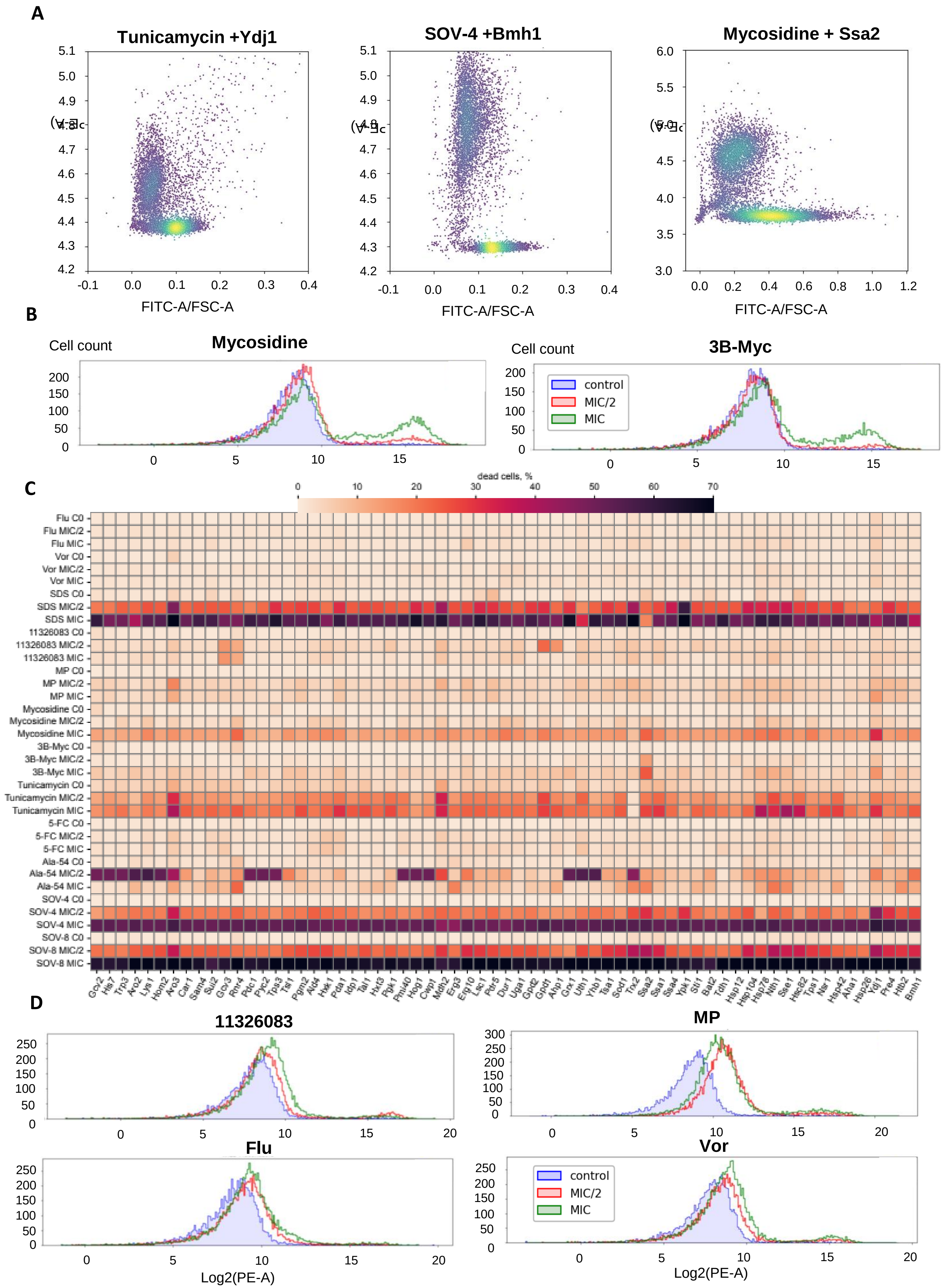


**Figure S2. (A) Flow cytometric analysis of protein leakage from dead cells (B) standard fluorescence of dead cells (C) the dead cell heatmap (D) weak staining with PI induced by several compounds**

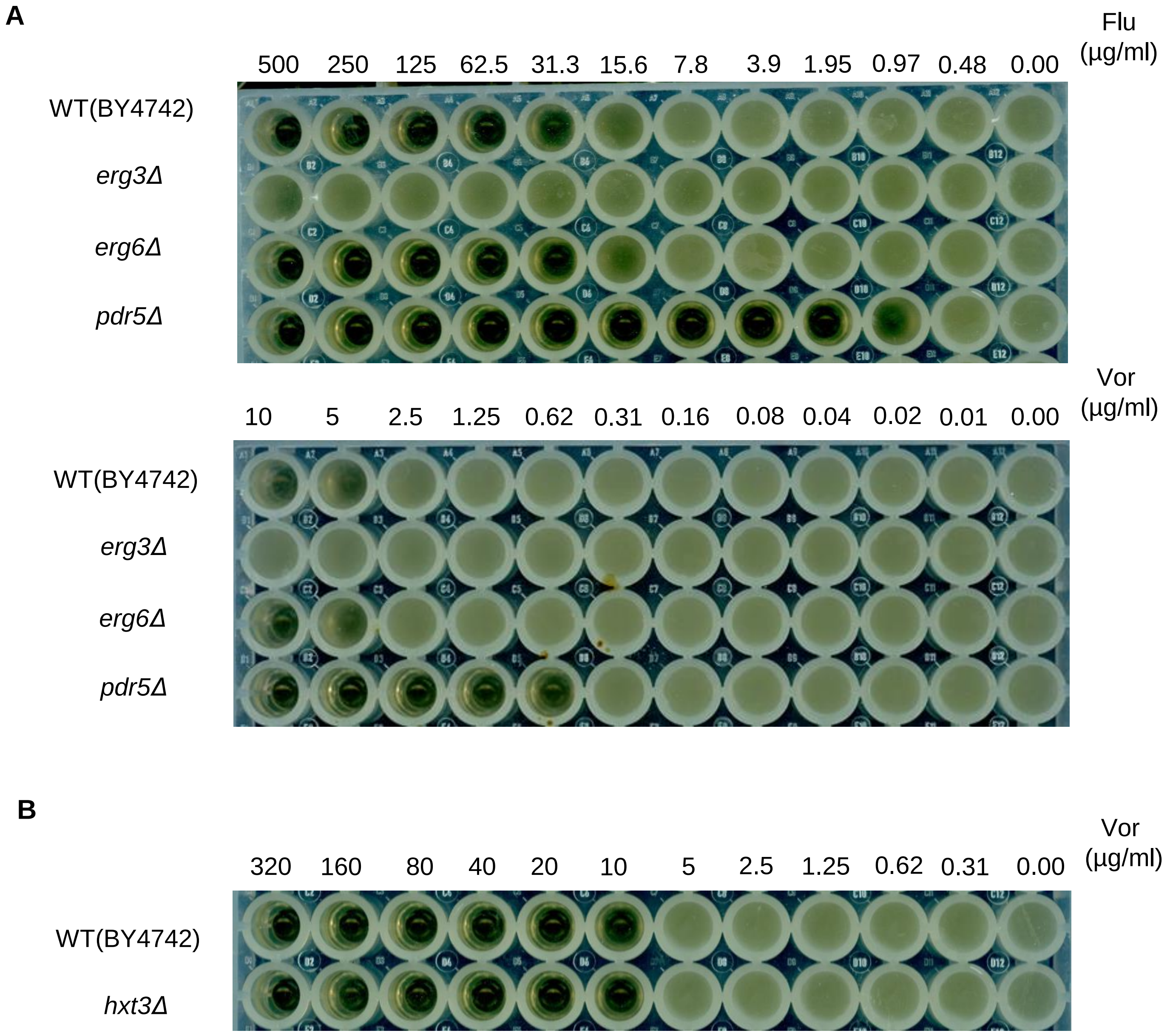


**Figure S3 Deletion of *ERG3* causes resistance to azoles, deletion of *PDR5* increases sensitivity, while deletion of *ERG6* and *HXT3* have little effect.** (A) Effect of the noted gene deletions on the MIC of the noted compounds; (B) Effect of the noted gene deletions on the MIC of the noted compounds.

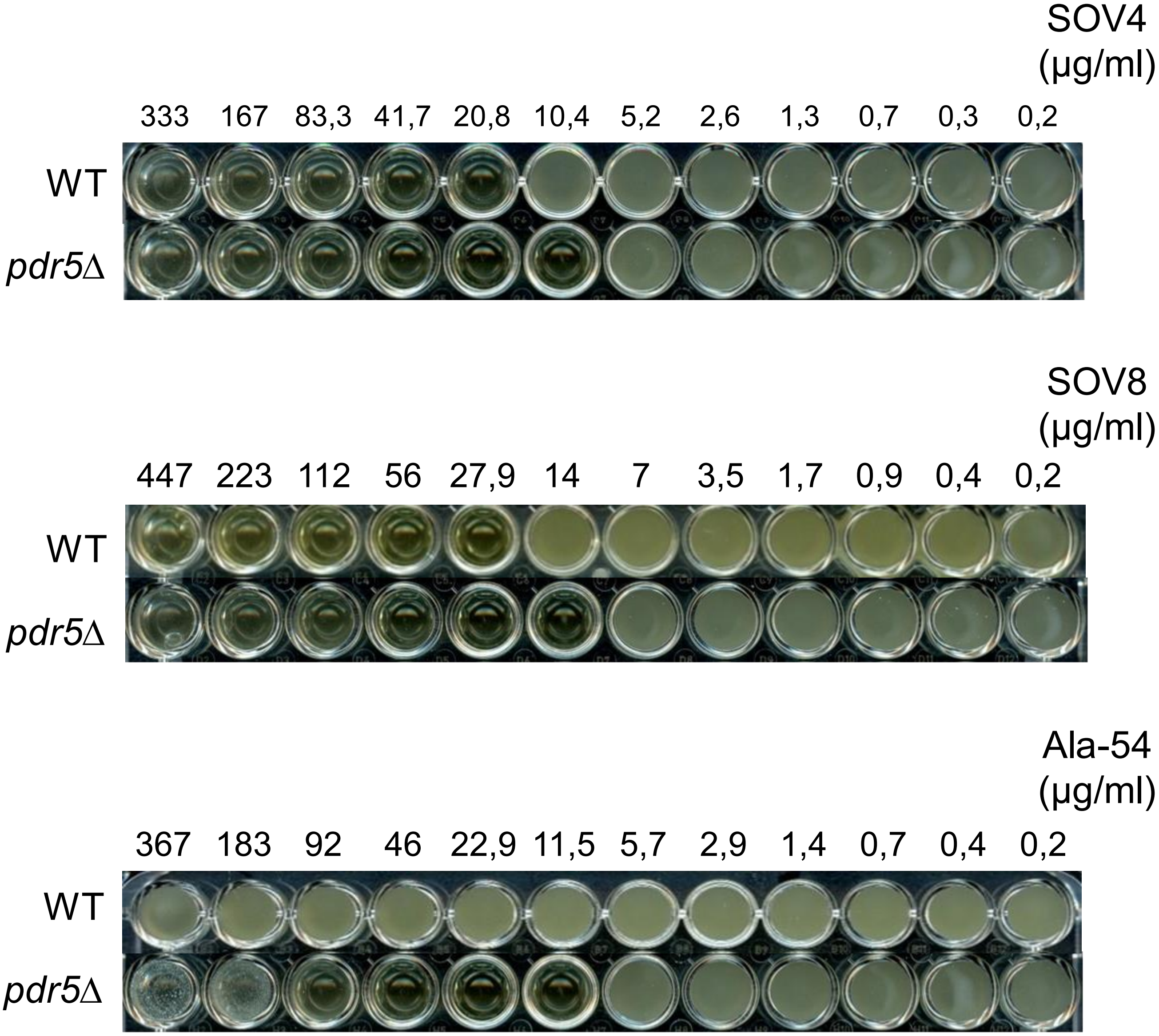


**Figure S4A. *PDR5* deletion has no effect on SOV4 and SOV8 activity, but Ala-54 becomes nearly as toxic as the other derivatives only after Pdr5 deletion.** Effect of the noted gene deletions on the MIC of the noted compounds.

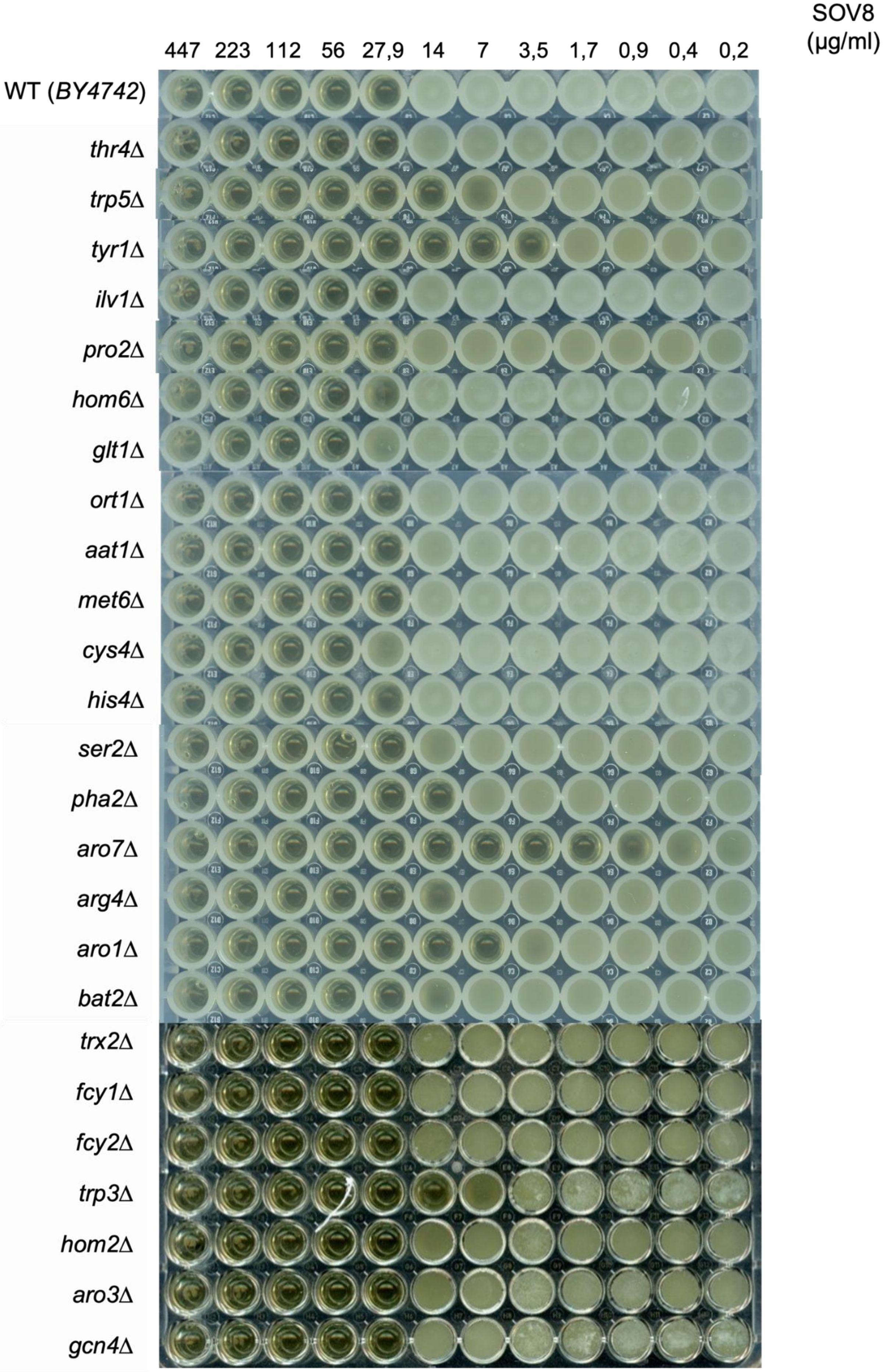


**Figure S4B. Deletion of tyrosine and tryptophan biosynthesis genes cause increasing of sensitivity to SOV4, while perturbation of biosynthetic pathways of other amino acids has no affect.** Effect of the noted gene deletions on the MIC of the noted compounds.

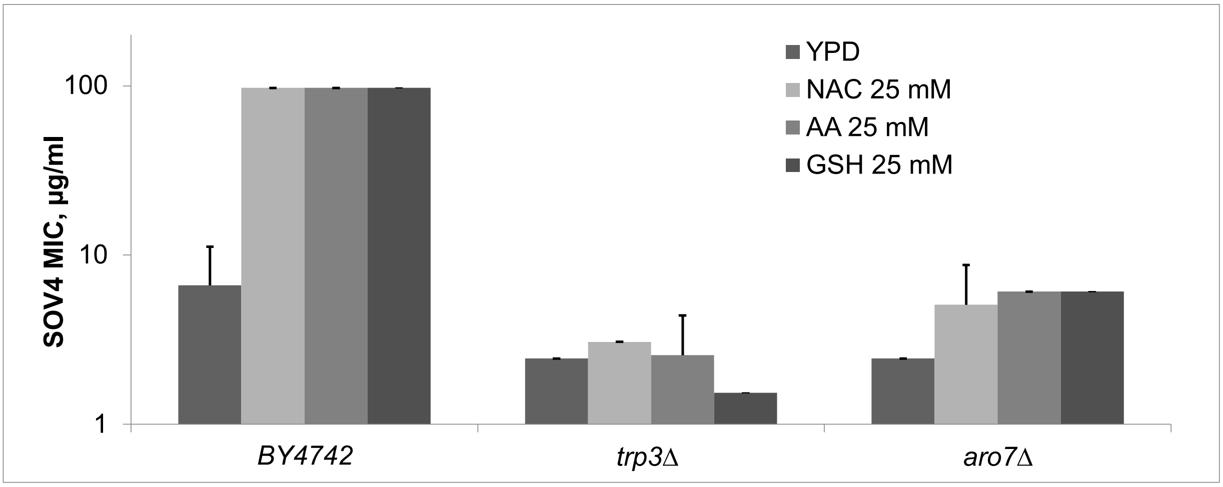


**Figure S4C. Sensitivity of the wild-type strain could be reduced by addition of antioxidants N-acetyl cysteine, ascorbic acid and glutathione, but aromatic amino acid biosynthesis mutants cannot be protected by antioxidant treatment.**
